# Structural basis of BRD4 association with P-TEFb

**DOI:** 10.64898/2026.09.03.749171

**Authors:** Abdallah A. Mohamed, Seychelle M. Vos

## Abstract

Cyclin-dependent kinases (CDKs) dynamically regulate RNA Polymerase (Pol) II during each stage of transcription. Phosphorylation of RNA Pol II and associated transcription factors by CDKs enables stage-specific exchange of factors. CDK9 and a cyclin partner (CYCT1, T2 or K) comprise the Positive Transcription Elongation Factor-b (P-TEFb) complex, which is responsible for releasing RNA Pol II from promoter proximal pausing into processive elongation. P-TEFb activity is stimulated by bromodomain-containing Protein 4 (BRD4); however, the structural basis for BRD4 association with P-TEFb has remained unclear. Using cryogenic-electron microscopy, AlphaFold modeling, and biochemical probing, we show that the P-TEFb interacting domain of BRD4 (BRD4-PID) interacts with P-TEFb via the C-lobe of CDK9 and the second cyclin box of CYCT1. The BRD4 binding surface on CDK9 is also used by super elongation complex component AFF4. Our analysis indicates that BRD4 may stimulate P-TEFb activity by remodeling the kinase αC helix relative to previous structures. Finally, P-TEFb activity is restricted when bound to the 7SK RNP complex. We show that in addition to BRD4-PID, the AFF4 N-terminus can release P-TEFb from the 7SK RNP complex. Together, these findings define how P-TEFb activity is regulated by its incorporation into distinct complexes.

**SIGNIFICANCE STATEMENT:** The exquisite control of gene expression is required for proper development and cellular response to environmental stress. Positive transcription elongation factor b (P-TEFb) is required for the phosphorylation of RNA Pol II and leads to the coordinated release of RNA Pol II from pause sites near the promoter. A subset of P-TEFb in the nucleus is bound to bromo-domain containing protein 4 (BRD4), which stimulates P-TEFb activity; however, it is unknown how BRD4 associates with P-TEFb. Here we report a cryo-EM structure of P-TEFb bound to BRD4. By comparing our structure with previously reported structures, we show that BRD4 and AFF4 bind to P-TEFb using an overlapping binding surface. Additionally, we observe that AFF4 relieves HEXIM1 mediated repression of P-TEFb.

## INTRODUCTION

RNA Polymerase (Pol) II transcribes most eukaryotic protein coding genes and is highly regulated to ensure appropriate gene expression.^1–3^ The association of distinct protein factors with RNA Pol II across the gene body modulates RNA Pol II activity.^1,3,4^ This differential factor association is partially regulated by the phosphorylation status of the largest RNA Pol II subunit, RPB1.^5,6^ RPB1 contains a C-terminal domain (CTD) composed of the heptapeptide repeat Y_1_S_2_P_3_T_4_S_5_P_6_S_7_.^1,5^ During transcription initiation, the initially hypophosphorylated CTD is modified by the kinase CDK7 to release initiation factors like Mediator from RNA Pol II and to allow for initial transcription.^7–9^ After promoter escape, metazoan RNA Pol II frequently pauses 25–100 bp from the transcription start site.^1^ This pause is initially induced by the underlying DNA sequence and is further modulated by transcription factors DRB sensitivity-inducing factor (DSIF) and negative elongation factor (NELF).^10–14^ Pause release and the assembly of productive elongation complexes requires the kinase positive transcription elongation factor b (P-TEFb).^15^ P-TEFb phosphorylates DSIF and NELF, as well as the RPB1 CTD and the RPB1 CTD linker.^13,15–19^ The phosphorylation of RNA Pol II and associated factors by P-TEFb allows for the displacement of NELF by elongation factors such as PAF1c and SPT6 and the transition of RNA Pol II from a paused state into a productive elongation state.^20,21^

P-TEFb is a heterodimer, composed of cyclin dependent kinase 9 (CDK9), and its predominant cyclin binding partner, cyclin-T1 (CYCT1), although it can associate with other cyclins, like cyclin-T2 and cyclin-K.^5,22–24^ CDK kinase activity is activated by the binding of its cyclin subunit and the phosphorylation of its T-loop, a conserved feature across CDKs.^25,26^ When CDKs are not bound to their cyclin partner, the T-loop is bound within the active site of the CDK, preventing substrate loading.^26,27^ Upon partner cyclin binding, the T-loop is rearranged, freeing the active site.^26,27^ CDK9’s activity is further regulated by the phosphorylation of T-loop residue Thr186 through CDK9’s autophosphorylation activity or by CDK-Activating Kinase (CAK), which is composed of CDK7, Cyclin H, and MAT1.^28,29^

P-TEFb activity is also regulated by its association with specific inactivating or activating protein binding partners. P-TEFb activity is inhibited when it is sequestered in the 7SK ribonucleoprotein (RNP) complex.^30,31^ Specifically, the HEXIM1 subunit of the 7SK RNP inhibits P-TEFb activity by associating with the CDK9 active site via its PYNT motif and inhibition is primarily mediated through residue Thr205 of the motif.^32–34^ Additional binding interfaces are formed between CYCT1 and the C-terminal HEXIM1 dimerization interface.^35^ In contrast, active P-TEFb containing complexes include the super elongation complex (SEC) or the bromodomain containing protein 4 (BRD4)–P-TEFb complex.^36–41^ The N-termini of SEC subunits AFF1/4 associate with P-TEFb.^42,43^ In addition to regulating transcription in uninfected cells, the SEC is involved in HIV-1 transcription, where AFF1/4 and HIV-1 Tat bind P-TEFb simultaneously.^22,40–42^ Structural and biochemical work have shown that the P-TEFb–AFF4–Tat complex enhances the association of HIV-1 Tat with the HIV-1 Transactivation Response element (TAR) RNA hairpin associated with paused RNA Pol II.^43– 46^ This interaction localizes P-TEFb to paused RNA Pol II to facilitate productive HIV-1 transcription.^41,44,47,48^

Human BRD4 is a 1362 amino acid protein that contains two N-terminal bromodomains that bind to acetylated histones, and an extra terminal (ET) domain that binds to other protein binding partners, such as JMJD6.^49–53^ The last 63 amino acids of the largely unstructured BRD4 C-terminus bind to P-TEFb and are referred to as the P-TEFb interaction domain (BRD4-PID).^54^ BRD4-PID can compete with the 7SK RNP complex for P-TEFb binding and stimulate P-TEFb’s kinase activity.^36–38,55^ Recent work has shown that the BRD4-PID is necessary for RNA Pol II to enter the gene body, suggesting that BRD4-PID is not only important for P-TEFb stimulation, but also important for transcription elongation.^56^ Biochemical work has been conducted to understand how BRD4-PID stimulates P-TEFb, identifying key motifs in BRD4-PID, namely BRD4-PID’s arginine rich motif (ARM) and leucine motif, that are important for the stimulation of P-TEFb activity.^55^ Finally, BRD4 and Tat have been reported to compete for P-TEFb binding, and inhibition of BRD4 by the bromodomain binding small molecule JQ1 leads to the transactivation of HIV-1.^54,57^ How HIV-1 Tat and BRD4-PID compete for P-TEFb association is not well understood.

We currently lack a structural understanding of how P-TEFb interacts with BRD4-PID and therefore the precise mechanism of how BRD4-PID activates P-TEFb. Furthermore, it is unknown how BRD4 releases P-TEFb from HEXIM1. To address these questions, we have taken a structural and biochemical approach to characterize the P-TEFb–BRD4-PID complex. Our results indicate that P-TEFb uses an overlapping binding surface to interact with either BRD4 or AFF4 and show that both BRD4 and AFF4 can release P-TEFb from HEXIM1 inhibition. Together, these results provide a framework to understand how P-TEFb activity is regulated in distinct protein complexes.

## RESULTS AND DISCUSSION

### Biochemical characterization of the P-TEFb–BRD4-PID complex

To investigate how bromodomain containing protein 4 (BRD4) binds and stimulates positive transcription elongation factor b (P-TEFb), we recombinantly co-expressed and purified a human P-TEFb variant containing full length human CDK9 and a stable CYCT1 truncation (amino acids 1–272). We separately isolated the human BRD4 P-TEFb Interacting Domain (BRD4-PID) (amino acids 1300–1362) **(Figures 1A and S1A)**. BRD4-PID and P-TEFb stably interacted in analytical size exclusion chromatography experiments **(Figure S1B)**. We next determined the binding affinity of BRD4-PID for P-TEFb using bio-layer interferometry (BLI) and a biotinylated version of the BRD4-PID. The measured affinity of BRD4-PID for P-TEFb is a K_d_ of 100 nM ± 30 nM **(Figure 1B)**. Our results differ from a previous publication where an affinity of 470 ± 140 nM was reported.^55^ The previous experiments were performed in a buffer containing 250 mM NaCl, whereas our buffer contained 100 mM NaCl. Salt sensitive association has been noted in prior co-immunoprecipitation experiments where stable P-TEFb–BRD4-PID interactions were only detected in buffers containing 200 mM or lower KCl concentrations.^36^

**Figure 1.**
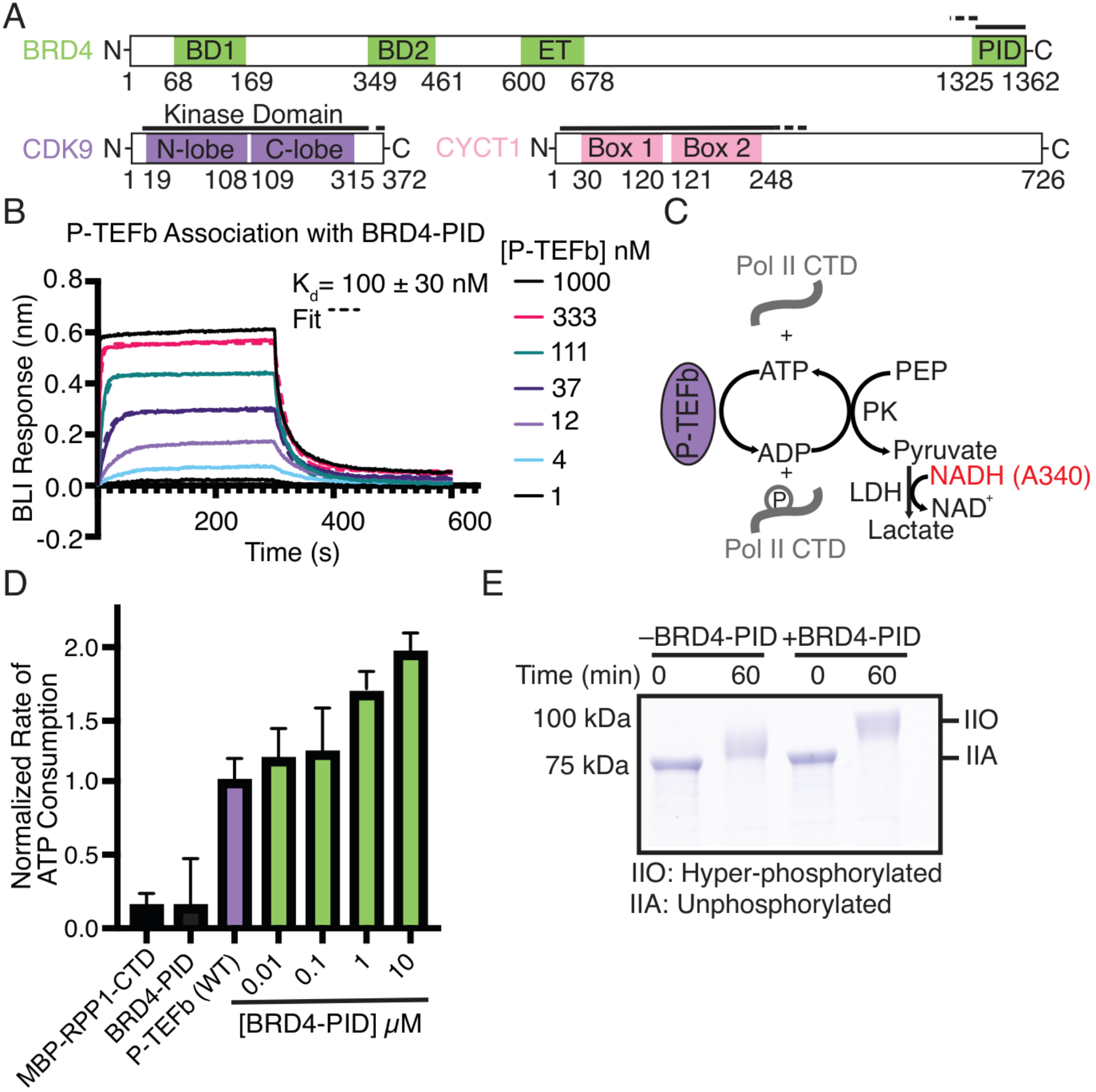
BRD4-PID binds and stimulates P-TEFb activity. (A) Domain architecture of BRD4 (green), CDK9 (purple), and CYCT1 (pink) with residue numbers indicating domain boundaries. Solid lines indicate the regions of the protein modelled in cryo-EM reconstructions reported here. Dashed lines indicate the regions of protein present in protein constructs but were not modeled. (B) P-TEFb binding to BRD4-PID as detected by Bio-layer interferometry (BLI). Concentration ranges shown. K_d_ was calculated from concentrations 333 nM, 111 nM, and 37 nM. Dashed lines indicate the fit used to calculate the K_d_. K_d_ was calculated from an average of three replicates and is 100 nM ± 30 nM. (C) Schematic of modified NADH-coupled ATPase assay used to test P-TEFb kinase activity. (D) P-TEFb kinase activity measured using the NADH-coupled ATPase assay. Bars correspond to the normalized change in absorbance at 340 nm as a function of time relative to P-TEFb kinase activity in the presence of MBP-RPB1-CTD. Each bar corresponds to the mean of three individual experiments. BRD4-PID titration shown, as well as the background signal from MBP-RPB1-CTD and BRD4-PID. Error bars represent the standard deviation between three individual experiments. (E) Gel based assay to detect phosphorylation of MBP-RPB1-CTD by SDS-PAGE. P-TEFb was incubated with and without BRD4-PID for one hour. Addition of phospho-groups onto MBP-RPB1-CTD cause the protein to migrate slower in SDS-PAGE due to the increase in negative charges. The positions of unphosphorylated (IIA) and hyper-phosphorylated (IIO) MBP-RPB1-CTD are indicated.

We next tested whether recombinantly purified BRD4-PID stimulated P-TEFb kinase activity. P-TEFb was incubated with increasing amounts of BRD4-PID in the presence of recombinantly expressed and purified human RPB1-CTD (amino acids 1593–1970) fused to maltose binding protein (MBP) and activity was monitored using a modified NADH-coupled ATPase assay **(Figure 1C)**. Under our assay conditions, it is not feasible to regenerate the substrate (MBP-RPB1-CTD) meaning the assay was conducted under substrate depleting conditions **(Figure 1C)**. Minimal background signal was observed from the MBP-RPB1-CTD on its own and the MBP-RPB1-CTD in the presence of BRD4-PID, indicating that ATPase activity is derived from P-TEFb’s kinase activity **(Figure 1D)**. BRD4-PID stimulated P-TEFb ATPase activity in a titratable fashion as previously observed, resulting in a nearly two-fold increase in ATPase activity at the maximum concentration of BRD4-PID tested **(Figure 1D)**.^55^

P-TEFb kinase activity can also be monitored by SDS-PAGE.^55^ The migration of hyper-phosphorylated MBP-RPB1-CTD (IIO) is retarded relative to unphosphorylated RPB1 CTD (IIA) in SDS-PAGE due to the presence of multiple negatively charged phosphate groups. Like the ATPase assays, P-TEFb incubated in the presence of a 100-fold molar excess of BRD4-PID produced a hyper-phosphorlyated CTD faster than isolated P-TEFb **(Figures 1E and S1C)**. No detectable shift in MBP-RPB1-CTD was observed when the MBP-RPB1-CTD was only incubated with BRD4-PID **(Figure S1D)**. Together, these results show that purified BRD4-PID associates with P-TEFb and stimulates P-TEFb kinase activity.

### Structure of the P-TEFb–BRD4-PID complex

We next visualized the P-TEFb–BRD4-PID complex using single particle cryo-EM. Purified P-TEFb and BRD4-PID along with ATPγS were incubated, applied to cryo-EM grids, and subjected to plunge freezing without crosslinking. Initial data processing indicated orientation bias in the sample. Thus, a subsequent data collection was conducted at a tilt angle of 15º. The tilted and non-tilted data sets were separately processed through particle picking and multiple rounds of 2D classification. Particles found in 2D classes with high resolution information and multiple views of the P-TEFb–BRD4-PID complex were selected, and the two particle sets were combined. *Ab-initio* classification was performed on the curated particle set and resulted in an initial map with density for P-TEFb and the BRD4-PID. Further classification and refinement resulted in a map that had a nominal resolution of 4.3 Å (gold standard FSC 0.143) **(Figures 2, S2, S3, and Supplementary Information, Table S1)**. The model includes amino acids 1318–1358 of BRD4-PID, amino acids 17–321 of CDK9, and amino acids 10–248 of CYCT1, and density for ATPγS in the active site **(Figures 1A, S3)**. The model shows good stereochemistry **(Supplementary Information, Table S1)**.

**Figure 2.**
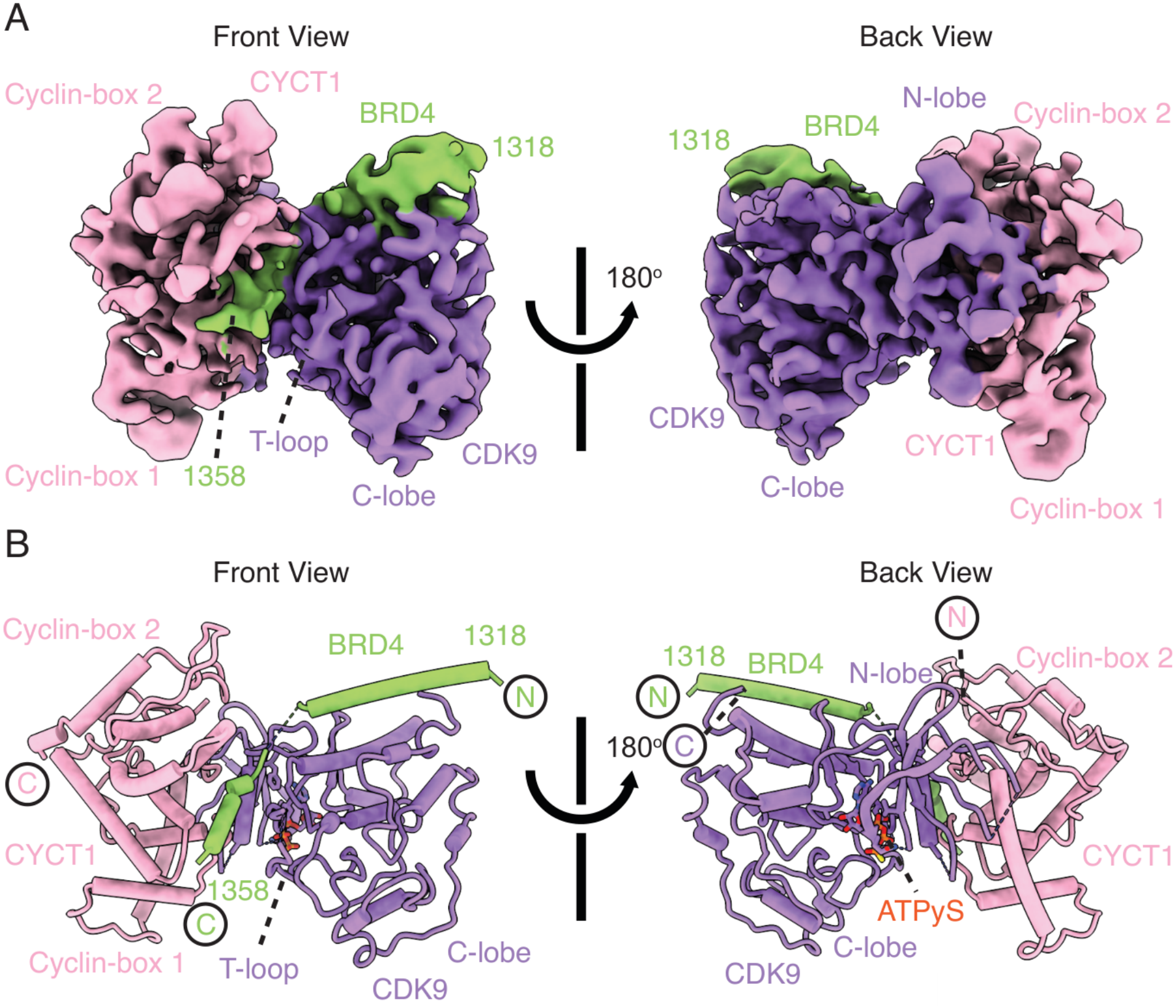
Structure of the P-TEFb–BRD4-PID complex. (A) Cryo-EM density map of the P-TEFb–BRD4-PID complex. BRD4 green, CYCT1 pink, CDK9 purple. Front and back views are shown. (B) Structure of the P-TEFb–BRD4-PID complex. Same coloring and views as in (A).

Our model shows that the BRD4-PID binds to both CDK9 and CYCT1 via two α helices. BRD4-PID α-helix 1 (residues 1318–1342) interacts with the C-terminal lobe of CDK9 while α-helix 2 (residues 1349– 1358) is sandwiched in between the CDK9 T-loop and CYCT1 and appears to contact the second cyclin box of CYCT1 **(Figures 1A and 2)**. Comparing our model to that of P-TEFb bound to ATP (PDB ID: 3BLQ), we observe that the buried surface area between CDK9 and CYCT1 increased by 1.2-fold upon the binding of BRD4-PID, from an area of 909.3 Å^2^ in the ATP bound state to an area of 1059.7 Å^2^ in the BRD4-PID bound state. These observations are consistent with previous experiments that showed robust BRD4-PID association with the P-TEFb complex but not with CYCT1 on its own.^55^

### Structural comparisons between P-TEFb–BRD4-PID and other P-TEFb complexes

We next compared our P-TEFb–BRD4-PID model to previously solved X-ray crystal structures of P-TEFb bound to ATP or other protein factors. Each crystal structure was determined in a different space group, and thus crystal packing constraints could influence observed differences between structures. All P-TEFb structures solved to date are in an active kinase conformation. The P-TEFb–BRD4-PID model is also in an active conformation but is contracted relative to all other structures **(Figure 3 and S4A–D)**.^29^ Specifically, BRD4-PID leads to the repositioning of the second cyclin box in CYCT1, which is relatively immobile in all other reported structures **(Figure 3A–B)**. The distance between CDK9 and the second cyclin box of CYCT1 is 22.6 Å in an ATP bound structure of P-TEFb (PDB ID: 3BLQ) and is reduced to 18.5 Å in the presence of BRD4-PID **(Figures S4A and S4D)**. Structures of HIV-1 Tat or AFF4-Tat bound P-TEFb retain the ~23 Å distance between CDK9 and CYCT1 as observed in the ATP bound structure **(Figure S4B–D)**.^44,58^

**Figure 3.**
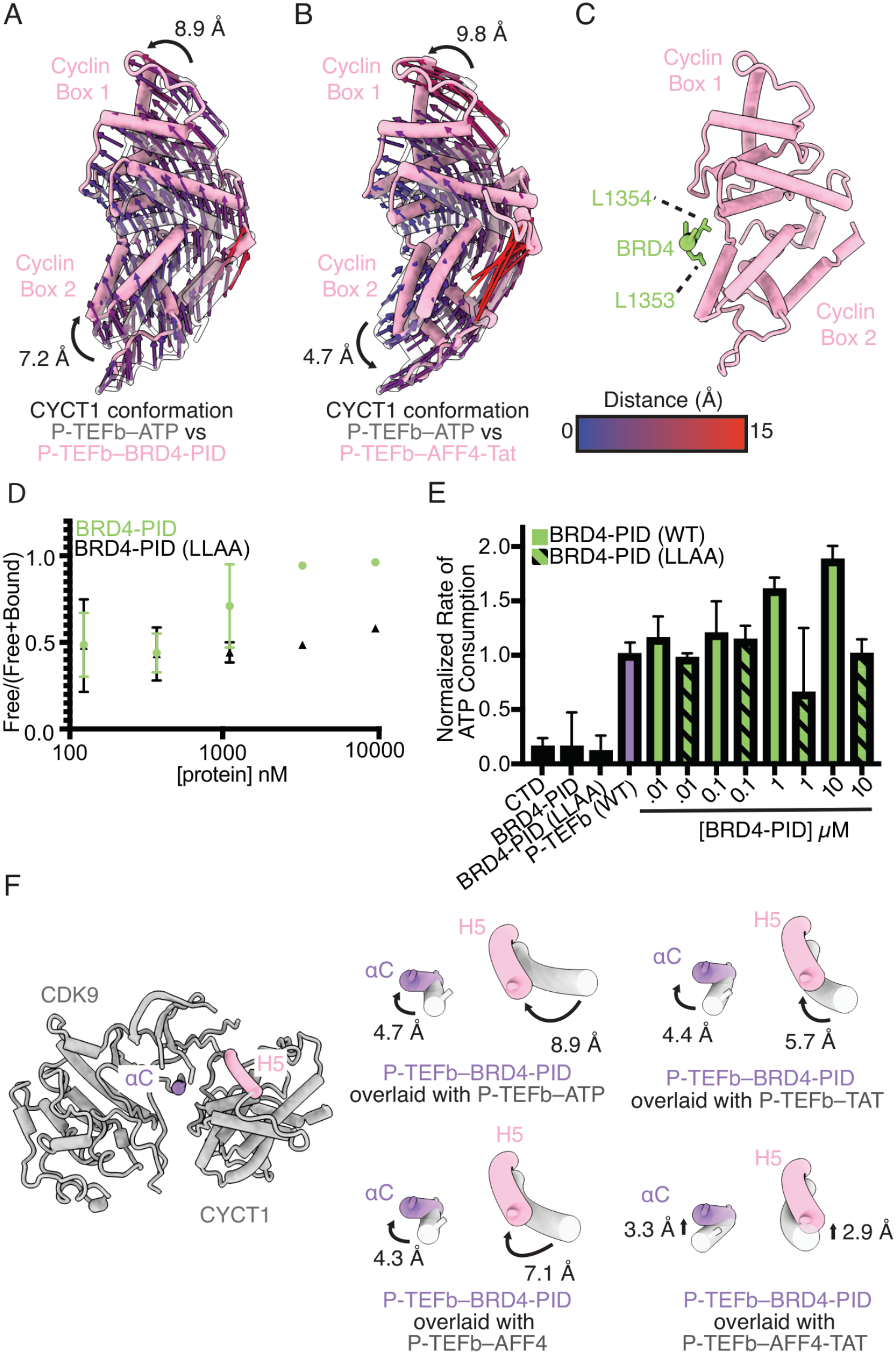
Structural rearrangements upon transcription factor binding to P-TEFb. (A) Overlay of CYCT1 when P-TEFb bound to BRD4 (pink) or ATP (grey) (PDB ID: 3BLQ). Structures aligned on CDK9 and rotated 120^°^ along the x-axis from the back view. Trajectories show the relative difference in amino acid positioning from the ATP bound state to the BRD4-PID bound state. Trajectories colored based on distance: blue (0 Å) to red (15 Å). Distances calculated for CYCT1 residues 124 and 163. (B) Overlay of CYCT1 when P-TEFb is bound to AFF4–Tat (pink) (PDB ID: 4OGR) or ATP (grey) (PDB ID: 3BLQ). Views and coloring the same as in (A). (C) Leucine 1353 and 1354 interact with CYCT1. Views and coloring the same as in (A). (D) Quantification of competition binding assay by WT BRD4-PID (green) or BRD4-PID (LLAA) (black). Error bars represent the standard deviation between three individual experiments. (E) P-TEFb kinase activity measured using the NADH-coupled ATPase assay as shown in Figure 1D with BRD4-PID (LLAA). (F) CDK9 αC and CYCT1 H5 rearrangements. Left: Position of the CDK9 αC (purple) and CYCT1 H5 (pink) relative to P-TEFb–BRD4-PID model (gray). Right: conformational changes for the αC (purple) and CYCT1 H5 (pink) elements in the BRD4-PID bound state relative to P-TEFb bound to ATP (PDB ID: 3BLQ), P-TEFb–Tat (PDB ID: 3MIA), P-TEFb–AFF4 (4IMY), and P-TEFb–AFF4–Tat (4OGR), all colored in grey. Distances measured from CDK9 amino acid 61 and CYCT1 amino acid 124, respectively.

The repositioning of the second cyclin box is likely caused by the association of BRD4 α2 with CYCT1 **(Figure 3A–B)**. Our structural model suggests that BRD4 residues Leu1353 and Leu1354 could coordinate binding to CYCT1 and likely contribute to the large-scale conformational changes we observe in cyclin box 2 **(Figure 3C)**. To test this idea, we generated a BRD4-PID construct substituting BRD4-PID LeuLeu1353– 1354AlaAla (LLAA). The relative affinity of LLAA for P-TEFb was assessed using a competition assay against Alexa Fluor 555 labeled WT BRD4-PID. Increasing concentrations of unlabeled WT BRD4-PID readily displaced labeled BRD4-PID from P-TEFb (**Figures 3D and S4E)** whereas LLAA weakly displaced labeled BRD4-PID **(Figures 3D and S4E)**. This indicates that LLAA binds to P-TEFb negligibly compared to WT BRD4-PID. Accordingly, LLAA does not stimulate P-TEFb ATPase activity, which is in line with published results using the LLAA BRD4-PID construct **(Figure 3E)**.^55^

The αC kinase element undergoes massive conformational changes during kinase activation to facilitate formation of the kinase active site.^59^ The CDK9 αC is in an active conformation in all published structures, and its motion is coordinated with CYCT1 α helix H5.^29,43–45,58^ Protein factors HIV-1 Tat (PDB ID: 3MIA), AFF4 (PDB ID: 4IMY), or AFF4 and Tat (PDB ID: 4OGR) result in a shift in CYCT1 α helix H5, and a slight rotation in the CDK9 αC helix and/or the loop between β3 and αC to various degrees relative to P-TEFb bound to ATP (PDB ID: 3BLQ) **(Figure 3F and S4F–I)**. BRD4-PID appears to exaggerate the changes in both elements. BRD4-PID appears to cause the CDK9 αC helix to move between 2–5 Å and the CYCT1 α helix H5 between 2–9 Å, depending on whether other protein factors, such as AFF4 or HIV-1 Tat, are bound to P-TEFb. In summary, BRD4 association with P-TEFb leads to more extensive conformational changes than observed in prior structures of P-TEFb.

### BRD4-PID and HIV-1 Tat bind on an overlapping surface on P-TEFb

Expression of HIV-1 is dependent on the recruitment of P-TEFb via HIV-1 Tat to the HIV-1 TAR RNA. BRD4-PID and HIV-1 Tat both stimulate P-TEFb activity and share sequence homology, including arginine rich and leucine motifs.^43,55^ Prior work showed that BRD4 residues 1209–1362 but not residues 1209–1328 displaced HIV-1 Tat from binding P-TEFb.^54^ Additionally, overexpression of BRD4-PID led to reduced expression of HIV-1 in J-Lat clone A2 cells.^54^ These findings suggested there may be a direct competition between BRD4 and HIV-1 Tat for P-TEFb.^54^ To examine this possibility, we compared our structure to a structure of P-TEFb bound to HIV-1 Tat.^58^ This analysis showed that BRD4 and HIV-1 Tat bound an overlapping binding surface on CYCT1. Specifically, an N-terminal loop in HIV-1 Tat (residues 7–13) clashes with BRD4-PID α2, providing an explanation for the previous biochemical and cell biological results **(Figure S4J)**.

### BRD4-PID and HPV E2 bind on an overlapping surface on P-TEFb

In addition to P-TEFb, BRD4-PID binds the human papillomavirus (HPV) E2 protein.^60,61^ The E2 protein among other roles is responsible for non-covalently tethering the papillomavirus genome to the host genome and for activating and repressing HPV gene expression.^60,61^ An X-ray crystal structure of BRD4-PID with the E2 protein showed that BRD4-PID contacted the E2 protein using α2 (residues 1349–1362).^62^ Although the BRD4-PID construct used in the E2 co-crystal structure contained the residues that interact with CDK9 (1325–1348), these residues are not resolved. Overlaying our structure with the BRD4-E2 structure showed that BRD4 would not be able to interact with both E2 and P-TEFb simultaneously **(Figure S4K)**. This is in line with previous work that has shown that E2 binding to BRD4 inhibits the interaction between P-TEFb and BRD4.^63^ It is thought that this competition prevents the recruitment of P-TEFb to viral oncogenes E6/E7 of the integrated HPV genome to repress their expression.^63^

### P-TEFb binds BRD4 and AFF4 at a shared CDK9 interface using a conserved sequence motif

Active P-TEFb exists in at least two distinct complexes in the cell, those bound to BRD4 and those bound to the super elongation complex subunits AFF1/4. Published work indicates that BRD4 in complex with P-TEFb is required for proper basal transcription activation and in response to stresses such as hypoxia.^64^ AFF4 and the super elongation complex are associated with transcriptional activation in response to stress.^65^ Therefore, these different pools of active P-TEFb appear to regulate RNA polymerase II activity in a gene and context specific manner. To better understand these observations, we compared structures of P-TEFb bound to AFF4 with our P-TEFb–BRD4-PID structure. In all reported structures and in contrast to HIV-1 Tat and BRD4-PID, AFF4 does not come in proximity with CDK9’s T-loop. AFF4 additionally has no reported stimulatory function and may even slightly inhibit P-TEFb kinase activity.^43^

The N-terminal α0 of AFF4 interacts with the C-lobe of CDK9 **(Figure 4A)**.^43–45^ This AFF4–CDK9 interaction was not extensively studied because it was thought to be a crystallization artifact at the unit cell interface of the AFF4–P-TEFb crystal structure. Furthermore, alanine scanning mutagenesis of AFF4 α0 had less impact on transcription activation in cells than AFF4 residues that interact with CYCT1.^43–45^ Notably, loss of the CDK9 binding interface reduced the affinity of AFF4 for P-TEFb.^43^

**Figure 4.**
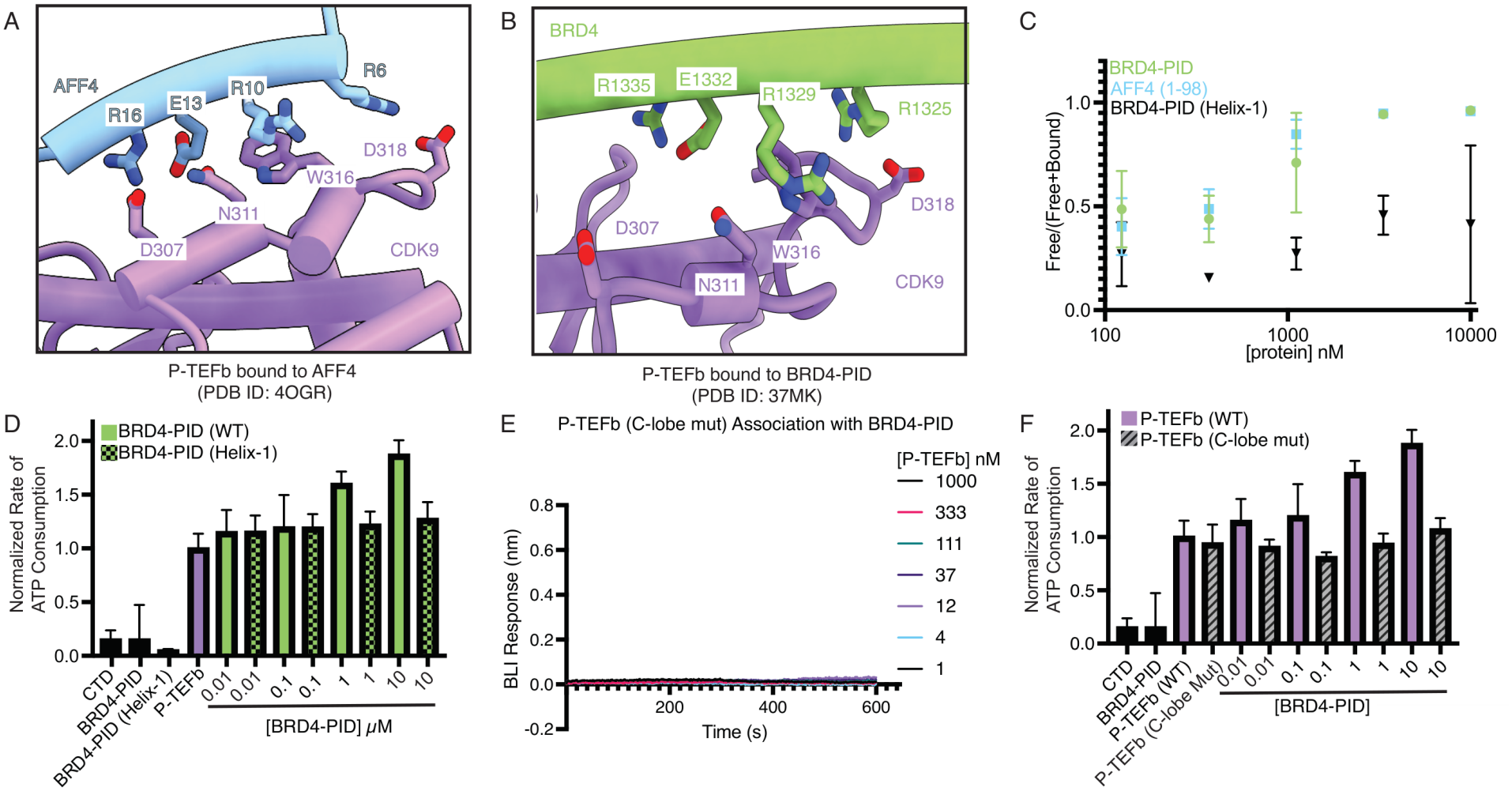
BRD4 and AFF4 interact with the C-lobe of P-TEFb. (A) Interaction between AFF4 (blue) and CDK9’s C-lobe (purple) (PDB ID: 4OGR) when bound to HIV-1 Tat. Key residues for the interaction are shown. (B) Interaction between BRD4 (green) and CDK9’s C-lobe (purple). Key residues for the interaction are shown. (C) Quantification of competition binding assay by WT BRD4-PID (green), AFF4 (1–98) (blue), or BRD4-PID (Helix-1 mutant) (black). Error bars represent the standard deviation between three individual experiments. (D) P-TEFb kinase activity measured using the NADH-coupled ATPase assay as shown in Figure 1D with the BRD4-PID (Helix-1) mutant. (E) P-TEFb (C-lobe mutant) binding to BRD4-PID as detected by Bio-layer interferometry (BLI). Concentration ranges shown. (F) P-TEFb kinase activity measured using the NADH-coupled ATPase assay as shown in Figure 1D with P-TEFb (C-lobe mutant).

We next identified residues that may facilitate the interaction between BRD4-PID or AFF4 α0 and the CDK9 C-lobe. The residues that AFF4 and BRD4 use to interact with the C-lobe of CDK9 share sequence homology, with a sequence identity of 44% and a sequence similarity of 75%, suggesting that AFF4 and BRD4 use similar binding interactions **(Figures 4A–B and S5A–B)**. Specifically, four residues on BRD4-PID Arg1325, Arg1329, Glu1332, and Arg1335, and their conserved counterparts on AFF4 Arg6, Arg10, Glu13, and Arg16, appear to coordinate binding to CDK9 residues Asp318, Trp316, Asn311, and Asp307, respectively **(Figures 4A–B and S5C)**. BRD4 Arg1329 and Arg1335 have been implicated as a part of a larger cluster of basic residues on BRD4-PID that are involved in the stimulation of P-TEFb activity.^55^

To test whether AFF4 and BRD4 compete for a shared binding surface on the C-lobe of CDK9, we performed a competition binding experiment with Alexa Fluor 555 labeled WT BRD4-PID, unlabeled AFF4 1– 98, and P-TEFb. Increasing concentrations of AFF4 1–98 led to the displacement of labeled BRD4-PID from P-TEFb (**Figure 4C and S4E)**. These data support the idea that BRD4 and AFF4 bind P-TEFb in a mutually exclusive manner **(Figure 4C and S4E)**.

We next characterized the BRD4-CDK9 C-lobe interaction surface. BRD4-PID interacting residues were substituted to Arg1325Glu, Arg1329Glu, Glu1332Arg, and Arg1335Ala (Helix-1 mutant). The Helix-1 mutant weakly competed Alexa Fluor 555 labeled WT BRD4-PID from P-TEFb, indicating a reduced affinity for P-TEFb than WT BRD4-PID **(Figure 4C and S4E)**. The Helix-1 mutant did not stimulate P-TEFb ATPase activity **(Figure 4D)**, consistent with the reduced binding interaction between BRD4 and P-TEFb.

We next performed the reciprocal experiment by mutating the interacting CDK9 residues to Asp318Ala, Trp316Val, Asn311Ala, and Asp307Ala (C-lobe mutant). BLI performed with the WT BRD4-PID and the C-lobe mutant showed no detectable binding of BRD4-PID **(Figure 4E)**. Consistent with this ablation of binding between BRD4-PID and the C-lobe mutant, BRD4-PID was no longer able to stimulate the C-lobe mutant’s ATPase activity **(Figure 4F)**. Importantly, the ATPase activity of the C-lobe mutant is like WT P-TEFb in the absence of BRD4-PID, indicating that the CDK9 C-lobe interface is required for BRD4-PID to bind to P-TEFb and not the general activity of P-TEFb.

### P-TEFb is released from the 7SK RNP complex by either BRD4-PID or AFF4

Up to 90% of P-TEFb in cells is found in the 7SK RNP complex, which inactivates P-TEFb kinase activity through interactions with 7SK protein HEXIM1/2.^30–32,66,67^ P-TEFb is released from the 7SK RNP complex by BRD4-PID and HIV-1 Tat *in vitro* and in cells.^55,68,69^ There are presently no experimentally derived structures of P-TEFb bound to HEXIM1.^35^ To understand how HIV-1 Tat, BRD4, or AFF4 bound P-TEFb may accommodate HEXIM1, we acquired an AlphaFold 3 model of a HEXIM1–P-TEFb dimer. The 7SK RNP complex contains two HEXIM1 molecules that are dimerized by their C-terminal coiled-coil domains, and each HEXIM1 can bind a P-TEFb molecule **(Figure S6)**.^35,70,71^ HEXIM1 is predicted to contact CDK9 via its PYNT motif, as expected from prior structural and biochemical work, and is additionally predicted to make extensive contacts with CYCT1. Interestingly, the contacts between CYCT1 and HEXIM1 described in prior NMR work are not observed in the predicted structure.^32,34,35^ AFF4 and BRD4 appear to bind in an overlapping fashion with HEXIM1 on P-TEFb by competing with a HEXIM1 α-helix spanning amino acids 257–281 **(Figure 5A–B)**.^35,70^ We note that the AFF4 overlap with HEXIM1 is minimal. Comparison of X-ray crystal structures of P-TEFb bound to both AFF4 and Tat (PDB ID: 4OGR) show that Tat shares a larger overlapping binding surface with HEXIM1 than AFF4, suggesting that AFF4 and Tat could work additively to release P-TEFb from HEXIM1.^44^

**Figure 5.**
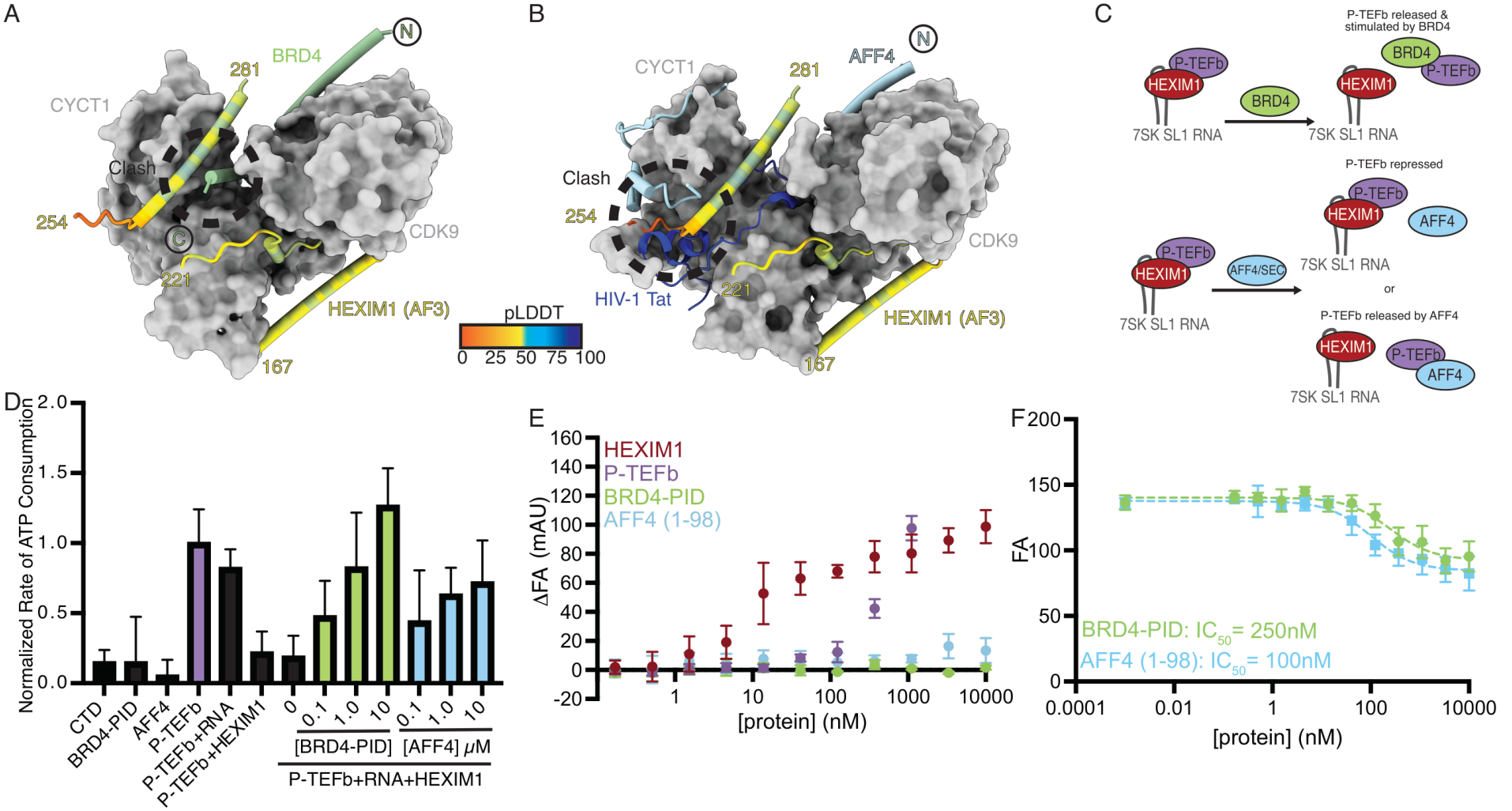
BRD4 and AFF4 release P-TEFb from the inhibited 7SK RNP complex. (A) AlphaFold 3 model of HEXIM1 bound to P-TEFb overlaid with our model of P-TEFb bound to BRD4-PID. P-TEFb shown as a gray surface. HEXIM1 colored according to pLDDT score (scale bar) and BRD4-PID shown in green. Location of HEXIM1 and BRD4-PID clash is shown. (B) AlphaFold 3 model of HEXIM1 bound to P-TEFb overlaid with the structure of P-TEFb bound to AFF4– Tat shown. P-TEFb shown as a gray surface. HEXIM1 colored according to pLDDT score (scale bar) and AFF4 shown in light blue and Tat blue. Location where HEXIM1 clashes with AFF4 and Tat are shown. (C) Cartoon illustrating the possible role that BRD4 and AFF4 play on P-TEFb activity in the context of the 7SK RNP complex. (D) P-TEFb kinase activity measured using the NADH-coupled ATPase assay. Bars correspond to the normalized change in absorbance at 340 nm as a function of time relative to P-TEFb kinase activity in the presence of MBP-RPB1-CTD. P-TEFb incubated with RNA and RPB1-CTD and P-TEFb incubated with HEXIM1 and RPB1-CTD shown. RPB1-CTD, as well as BRD4-PID and AFF4 (1–98) incubated with RPB1-CTD are also shown. P-TEFb incubated with RNA and HEXIM1 is shown. BRD4-PID and AFF4 (1–98) were titrated into the P-TEFb–HEXIM1–RNA complex. Error bars represent the standard deviation between three individual experiments. Each bar corresponds to the mean of three individual experiments. (E) Fluorescence anisotropy of the 7SK stem loop 1 RNA to HEXIM1, P-TEFb, BRD4-PID, and AFF4 (1–98). All data points were normalized to an RNA only control and displayed as ΔFA. Error bars correspond to the standard deviation between three individual experiments. (F) Competition fluorescence anisotropy assay where BRD4-PID or AFF4 (1–98) were titrated against a pre-formed P-TEFb–HEXIM1–7SK RNA complex. Dotted lines represent a curve fit. Error bars correspond to the standard deviation between three individual experiments. Calculated IC_50_ values included in panel.

From our structural prediction of HEXIM1 bound P-TEFb, we hypothesized that AFF4 may release P-TEFb from the 7SK RNP complex. To test this, we used a modified version of the NADH-coupled ATPase assay to measure P-TEFb activity in the context of the 7SK complex **(Figure 5C)**. Full-length HEXIM1 and 5’-FAM labeled stem loop 1 of the 7SK RNA (residues 26–85) that binds and releases an autoinhibitory patch on HEXIM1 were mixed with P-TEFb.^34,72^ Incubation of P-TEFb with the 7SK RNA on its own resulted in mild ATPase inhibition. In contrast, addition of HEXIM1 with or without stem loop 1 of the 7SK RNA and P-TEFb resulted in complete inhibition of P-TEFb’s apparent ATPase activity **(Figure 5D)**. Titration of BRD4-PID into reactions containing P-TEFb, HEXIM1, and the 7SK RNA reversed this inhibition while also stimulating P-TEFb in a titratable manner, whereas reactions lacking P-TEFb but containing all other protein factors showed no change in ATPase activity **(Figure 5D)**. This indicates that BRD4-PID activates P-TEFb, suggesting that BRD4-PID relieves P-TEFb inhibition by the 7SK RNP complex. We next conducted these experiments with the N-terminus of AFF4 (amino acids 1–98). Like BRD4, AFF4 addition resulted in a titratable increase in P-TEFb activity, but unlike BRD4, AFF4 did not lead to the stimulation of P-TEFb activity **(Figure 5D)**. Together these results show that both AFF4 and BRD4 can relieve P-TEFb ATPase inhibition that is mediated by HEXIM1 and the 7SK RNA, and BRD4-PID stimulates P-TEFb ATPase activity whereas AFF4 does not, as previously reported.^43,55^

Two models can explain how BRD4-PID and AFF4 relieve HEXIM1–7SK RNA induced inhibition of P-TEFb: (1) BRD4-PID and AFF4 remain bound to the inhibitory P-TEFb–HEXIM1–7SK RNA complex and their presence allows P-TEFb to adopt a catalytically productive conformation or (2) BRD4-PID and AFF4 displace P-TEFb from the inhibitory complex. To distinguish between these models, we conducted fluorescence anisotropy experiments. We first assessed how individual factors associated with the 5’-FAM labeled 7SK stem loop 1 RNA (HEXIM1, P-TEFb, AFF4 (1–98) and BRD4-PID). BRD4-PID and AFF4 displayed no detectable association with the 7SK stem loop 1 RNA **(Figure 5E)**. HEXIM1 strongly associated with 7SK stem loop 1 RNA, whereas P-TEFb only modestly associated with the RNA **(Figure 5E)**.

We next incubated the 7SK stem loop 1 RNA with fixed concentrations of P-TEFb and HEXIM1 and titrated in BRD4-PID and AFF4 (1–98). In both instances, the anisotropy signal decreased as increasing amounts of either BRD4-PID or AFF4 (1–98) were added. The resulting anisotropy signal at high BRD4-PID or AFF4 (1–98) concentrations corresponded to that observed for the HEXIM1–7SK complex (~80 mAU).

These results indicate that P-TEFb is removed from the 7SK–HEXIM1 complex **(Figure 5F)**. At a concentration of 300 nM HEXIM1 and P-TEFb, half-maximal inhibitory concentration (IC_50_) values of 250 nM for BRD4-PID and 100 nM for AFF4 (1–98) were determined. This is consistent with AFF4 binding P-TEFb more readily than BRD4.^43,55^ Our results suggest that the mechanism for P-TEFb release from the HEXIM1 complex could involve competition with an overlapping surface on CYCT1 between HEXIM1 and BRD4/AFF4, or by favoring a P-TEFb conformation that is not compatible for HEXIM1 binding. More work is required to understand the mechanism of P-TEFb release from HEXIM1 for both AFF4 and BRD4 **(Figure 5 and S6)**, including an experimentally derived structure of HEXIM1 bound to P-TEFb.

## CONCLUSIONS

Here we structurally and biochemically characterized the interaction between P-TEFb and BRD4. We additionally investigated the regulatory interplay between P-TEFb binding partners BRD4, AFF4, HEXIM1, and HIV-1 Tat. BRD4 appears to stimulate P-TEFb activity by stabilizing P-TEFb in a more active conformation. Additionally, BRD4 and AFF4 associate with the C-lobe of CDK9 using a similar sequence motif and surface, and, thus, do not appear to simultaneously bind P-TEFb. Finally, AFF4, like BRD4-PID and HIV-1 Tat, can release P-TEFb from the 7SK RNP complex. While we were in the final stages of preparing this manuscript, complementary results describing the interaction of BRD4-PID to CYCT1 and CYCT2 were reported using FragLite mapping.^73^ The authors identified BRD4-PID α helix-2 residues Leu1354 and Phe1357 as being important for BRD4-PID binding to CYCT2. These data along with our work show that BRD4 α helix-2 is an important feature of BRD4-PID that drives BRD4 interactions with P-TEFb.^73^

Proteins that associate with active P-TEFb can be classified into at least two categories: 1) those that stimulate P-TEFb’s kinase activity and release it from the 7SK RNP complex (*e*.*g*., HIV-1 Tat and BRD4), and 2) those that only release it from the 7SK RNP complex (e.g., AFF4). BRD4 and AFF4 differentially regulate gene expression.^64,65^ This separation of BRD4 and AFF4 function in cells could be linked to their differential influence on P-TEFb activity *in vitro*, where the fine-tuned expression of stress-response related genes requires a less active P-TEFb bound to AFF4, but basal transcription requires more active P-TEFb– BRD4. This remains to be explored further in future studies. It is unclear how different P-TEFb complexes could associate with RNA Pol II in a gene specific manner. Both the super elongation complex and BRD4 can associate with acetylated nucleosomes, and this could potentially facilitate P-TEFb recruitment to specific chromatin locations **(Figure 6)**.^49,74–76^ Further work is needed to understand how P-TEFb is incorporated into specific complexes and brought to paused RNA Pol II.

**Figure 6.**
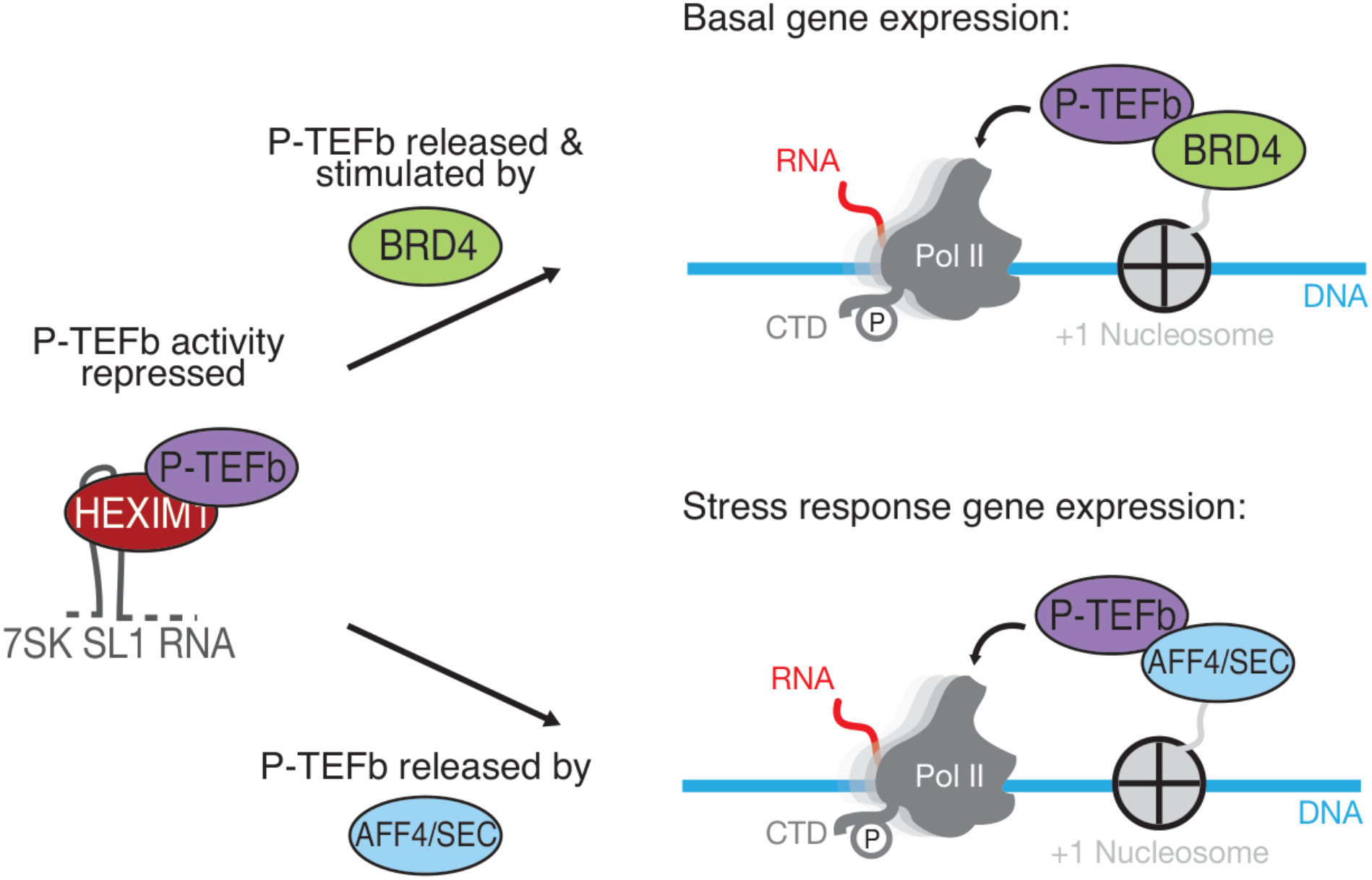
Model of P-TEFb role in transcription. P-TEFb activity is repressed when bound to HEXIM1 as a part of the 7SK RNP complex. BRD4 or AFF4 can release P-TEFb from the 7SK RNP complex, and recruit P-TEFb to RNA Pol II, leading to productive elongation.

## MATERIALS AND METHODS

Detailed materials and methods are provided in the accompanying Supplementary Information file.

### Cloning and Protein Expression

Human P-TEFb constructs were cloned into insect cell expression plasmids and expressed in Hi5 cells (Expression Systems). Human BRD4-PID and RPB1-CTD constructs were cloned into the 1C plasmid (Addgene: 29654) with an N-terminal His_6_-MBP tag followed by a TEV cleavage site and expressed in Bl21 (DE3) RIL cells. Human HEXIM1 and AFF4 (1–98) were cloned into the 1B plasmid with an N-terminal His_6_ tag followed by a TEV cleavage site (AddGene: 29653) and expressed in Rosetta 2 pLysS cells.

### Protein Purification

Proteins were purified using nickel and/or amylose affinity columns followed by size exclusion chromatography.

### Preparation of Alexa Fluor 555 labeled WT-BRD4 PID

The WT BRD4-PID construct with an N-terminal cystine was labeled using Alexa Fluor 555 C2-maleimide (ThermoFisher).

### Analytical Size-Exclusion Chromatography

P-TEFb was incubated with BRD4-PID and applied to a Superdex 200 3.2/300 column. Peak fractions were analyzed by SDS-PAGE.

### Bio-layer interferometry

Avi-tagged BRD4-PID were bound to Octet SA biosensors and incubated with P-TEFb (WT) or P-TEFb (C-lone mutant). Binding curves were fit for each replicate in GraphPad Prism Version 10 and the average and standard deviation of the K_d_ from each binding replicate were determined and reported.

### NADH-coupled ATPase Assay

A modified NADH coupled ATPase assay was used to measure relative rates of ATP hydrolysis of P-TEFb in the presence of BRD4-PID as previously described.^20,77^ For 7SK release assay, P-TEFb was incubated with HEXIM1 and 7SK RNA in the presence of either BRD4-PID or AFF4 (1–98).

### AlphaFold Modeling

AlphaFold 3 modeling was performed on the online AlphaFold 3 server for the P-TEFb–BRD4-PID and P-TEFb–HEXIM1 complexes.^78^

### Cryo-EM Sample Preparation

The P-TEFb–BRD4-PID complex was applied to R2/2 UltrAuFoil grids (Quantifoil) and vitrified using a a Vitrobot Mark IV (FEI Company).

### Cryo-EM Data Collection and Processing

Cryo-EM data were collected on a FEI Titan Krios with a K3 summit direct detector with a GIF quantum energy filter (Gatan). Two data collections were done, with a 0º and 15º tilt. Data sets were combined and processed in cryoSPARC v5.0.2 and Relion5.^79,80^

### Model Building

An AlphaFold 3 model of the P-TEFb–BRD4-PID complex was used for flexible fitting using ChimeraX v1.10.1, Namdinator, and Phenix v1.20.1-4487.^81–83^ The model was manually adjusted in Coot v1.3.1 and refined in Phenix v1.20.1-4487.^82,84^

### Multiple Sequence Alignment

Sequences for CDK9, BRD4, and AFF4 were obtained from UniProt. Sequences were aligned using MAFFT v7 online server and visualized in JalView v2.11.5.1.

### Fluorescence Anisotropy

A 5’ 6-FAM labeled 7SK RNA (26–85) was incubated with increasing concentrations of HEXIM1, P-TEFb, BRD4-PID or AFF4 (1–98). For competition assays, the 7SK RNA, HEXIM1, and P-TEFb were kept at a constant concentration and incubated with either BRD4-PID or AFF4 (1–98). Binding was measured using a Tecan Spark Plate reader.

## ACKNOWLEDGEMENTS

We thank past and present members of the Vos lab for helpful discussions. Cryo-EM specimens were prepared, and data were collected at the Cryo-EM facility at MIT.nano. We thank S. Sterling, D. Lim, and J. Podgorski for support at MIT.nano. We thank M. Yaffe for helpful discussions.

## AUTHOR CONTRIBUTIONS

A.A.M. and S.M.V. designed research; A.A.M. performed research; A.A.M. and S.M.V. contributed new reagents/analytic tools; A.A.M. and S.M.V. analyzed data; and A.A.M. and S.M.V. wrote the paper.

## DATA AND CODE AVAILABILITY

The cryo-EM map of the P-TEFb–BRD4-PID complex has been deposited in the Electron Microscopy Data Bank under accession code EMD-XXXXX, and the atomic coordinates have been deposited in the Protein Data Bank under accession code XXXX. All other data supporting the findings of this study are available within the article and its Supplementary Information.

