## Supplementary material for "Structural basis of BRD4 association with P-TEFb": Supplementary_Information_BRD4_P-TEFb.pdf

#### SUPPLEMENTARY MATERIALS AND METHODS

##### Cloning and Protein Expression

Human GST-P-TEFb and MBP-RPB1-CTD were cloned and expressed as previously described.[1]

Human MBP-P-TEFb and MBP-P-TEFb (C-lobe mutant) were cloned as follows: CDK9 and CDK9 (C-lobe mutant) plasmids were synthesized by Twist Bioscience. Full length CDK9 and CDK9 (C-lobe mutant) were inserted into a 438-A (AddGene: 55218) plasmid with an N-terminal TEV cleavable MBP tag. CYCT1 was inserted into a 438-A vector. CYCT1 and each respective CDK9 vector were combined into a pBig1a vector (AddGene: 80611) to produce the final P-TEFb (WT) and P-TEFb (C-lobe mutant) expression vector. The expression vectors were transformed into *E. coli* DH10EMBacY (Geneva Biotech) for bacmid generation. Positive colonies that contained the bacmid were isolated for amplification. Bacmids were isolated by alkaline lysis and ethanol precipitation. The bacmids were transfected into adherent Sf9 cells (Expression Systems) to produce V0 virus. V0 virus was amplified into V1 virus in Sf21 cells (Expression Systems). For large scale expressions, 1.2 L of Hi5 cells (Expression Systems) at a density of 1 million cells/mL were infected with V1 virus. After 72 hours cells were harvested by centrifugation and resuspended in lysis buffer (300 mM NaCl, 20 mM Na-HEPES pH 7.4, 10% (v/v) glycerol, 30 mM imidazole pH 8.0, and 1 mM DTT, 0.284 μg•mL$^{-1}$ leupeptin, 1.37 μg•mL$^{-1}$ pepstatin A, 0.17 mg•mL$^{-1}$ PMSF, and 0.33 mg•mL$^{-1}$ benzamidine). Resuspended cells were snap frozen in liquid nitrogen and stored at -80°C until purification.

Human BRD4-PID (BRD4 residues 1300–1362) was cloned into the 1C plasmid (Addgene: 29654) with an N-terminal MBP tag followed by a $His_6$ tag followed by a TEV cleavage site and an N-terminal tryptophan to aid in protein detection and quantification. The Avi-tagged BRD4-PID was cloned using sequence and ligation independent cloning (SLIC) by cloning the Avi tag after the TEV cleavage site. WT BRD4-PID and mutants LLAA and Helix-1 (LL1353–1354AA and R1325E R1329E E1332R R1335A) were purchased as gene fragments from Integrated DNA Technologies. The fragments contained an N-terminal Avi-tag followed by a cysteine residue and tryptophan and were cloned into vector 1C by Gibson assembly. All constructs were confirmed by full plasmid sequencing (Plasmidsaurus). The BRD4-PID constructs were transformed into Bl21 (DE3) RIL cells and grown at 37°C. Protein expression was induced with 0.5 mM IPTG when the cells reached an OD 600 nm of 0.6. Cells were grown for an additional 3 hours at 37°C. Cells were then harvested by centrifugation and resuspended in lysis buffer (300 mM NaCl, 20 mM Na-HEPES pH 7.4, 10% (v/v) glycerol, 30 mM imidazole pH 8.0, and 1 mM DTT, 0.284 μg•mL$^{-1}$ leupeptin, 1.37 μg•mL$^{-1}$ pepstatin A, 0.17 mg•mL$^{-1}$ PMSF, and 0.33 mg•mL$^{-1}$ benzamidine). Resuspended cells were snap frozen in liquid nitrogen and stored at -80°C until purification.

Full length human HEXIM1 and AFF4 (residues 1–98) were cloned into a 1B plasmid with an N-terminal $His_6$ tag followed by a TEV cleavage site (AddGene: 29653). The HEXIM1 plasmid was transformed into Rosetta 2 pLysS cells and grown at 37°C. Protein expression was induced with 0.5 mM IPTG when the cells reached an OD 600 nm of 0.6. Cells were grown for an additional 3 hours at 30°C. Cells were then harvested by centrifugation and resuspended in lysis buffer (300 mM NaCl, 20 mM Na-HEPES pH 7.4, 10% (v/v) glycerol, 30 mM imidazole pH 8.0, and 1 mM DTT, 0.284 μg mL$^{-1}$ leupeptin, 1.37 μg mL$^{-1}$ pepstatin A, 0.17 mg•mL$^{-1}$ PMSF, and 0.33 mg•mL$^{-1}$ benzamidine). Resuspended cells were snap frozen in liquid nitrogen and stored at -80°C until purification. The AFF4 plasmid was transformed into Rosetta 2 pLysS cells and grown at 37°C. Protein expression was induced with 0.5 mM IPTG when the cells reached an OD 600 nm of 0.8. Cells were grown for an additional 16 hours at 16°C. Cells were then harvested by centrifugation and resuspended in lysis buffer (500 mM NaCl, 20 mM Na-HEPES pH 7.4, 10% (v/v) glycerol, 30 mM imidazole pH 8.0, and 1 mM DTT, 0.284 μg•mL$^{-1}$ leupeptin, 1.37 μg•mL$^{-1}$ pepstatin A, 0.17 mg•mL$^{-1}$ PMSF, and 0.33 mg•mL$^{-1}$ benzamidine). Resuspended cells were snap frozen in liquid nitrogen and stored at -80°C until purification.

##### Protein Purification

All protein purification steps were performed at 4°C unless otherwise stated.

The GST tagged construct of P-TEFb, which was used for the cryo-EM sample, was purified from 2.8 liters of Hi5 expression. Cell pellets were thawed, lysed by sonication, and cleared by centrifugation. The clarified lysate was then filtered through a 5 µm syringe filter followed by a 0.45 µm syringe filter. The filtered lysate was applied to a 5 mL HisTrap HP column (Cytiva) that was equilibrated in lysis buffer (300 mM NaCl, 20 mM Na-HEPES pH 7.4, 10% (v/v) glycerol, 30 mM imidazole pH 8.0, and 1 mM DTT, 0.284 µg•mL$^{-1}$ leupeptin, 1.37 µg•mL$^{-1}$ pepstatin A, 0.17 mg•mL$^{-1}$ PMSF, and 0.33 mg•mL$^{-1}$ benzamidine). The HisTrap HP column was washed with ten column volumes of lysis buffer, followed by five column volumes of a high salt wash buffer (1000 mM NaCl, 20 mM Na-HEPES pH 7.4, 10% (v/v) glycerol, 30 mM imidazole pH 8.0, and 1 mM DTT) and five column volumes of 150 buffer (150 mM NaCl, 20 mM Na-HEPES pH 7.4, 10% (v/v) glycerol, 30 mM imidazole pH 8.0, and 1 mM DTT). A 5 mL HiTrap SP HP column (Cytiva) was equilibrated in nickel elution buffer (150 mM NaCl, 20 mM Na-HEPES pH 7.4, 10% (v/v) glycerol, 500 mM imidazole pH 8.0, and 1 mM DTT) and attached in-tandem with the HisTrap column. The protein was then eluted over a gradient from the 150-wash buffer to the nickel elution buffer over 30 minutes. After the elution concluded, the HisTrap column was removed and the HiTrap S column was washed with a S150 wash buffer (150 mM NaCl, 20 mM Na-HEPES pH 7.4, 10% (v/v) glycerol, and 1 mM DTT) for five column volumes. The protein was then eluted over a gradient from the S150 wash buffer with S1000 wash buffer (1000 mM NaCl, 20 mM Na-HEPES pH 7.4, 10% (v/v) glycerol, and 1 mM DTT) over 30 minutes. Peak fractions were pooled and mixed with 1.5 mg of His$_6$-Tabacco Etch Virus (TEV) protease and dialyzed overnight against lysis buffer.[2] The protein was then applied to a HisTrap equilibrated in lysis buffer to remove the His$_6$-TEV protease. Protein was concentrated in a 10 kDa MWCO Amicon Ultra Centrifugal Filter (Millipore) and applied to a HiLoad S200 16/600 pg column (Cytiva) equilibrated in SEC buffer (300 mM NaCl, 20 mM Na-HEPES pH 7.4, 10% (v/v) glycerol, and 1 mM DTT). Protein purity was assessed by SDS-PAGE followed by Coomassie staining. Peak fractions were concentrated in 10 kDa MWCO Amicon Ultra Centrifugal Filter (Millipore). The protein was stored in the SEC buffer. The protein was aliquoted, flash-frozen in liquid nitrogen, and stored at −80°C.

The MBP tagged P-TEFb (WT) and P-TEFb (C-lobe mutant), which were used for the biochemical assays, were purified as follows. 2.8 liters of Hi5 expression were used for each construct. Cell pellets were thawed, lysed by sonication, and cleared by centrifugation. The clarified lysate was then filtered through a 5 µm syringe filter followed by a 0.45 µm syringe filter. The filtered lysate was applied to a packed 10 mL amylose column (New England Biolabs) that was equilibrated in lysis buffer (300 mM NaCl, 20 mM Na-HEPES pH 7.4, 10% (v/v) glycerol, and 1 mM DTT, 0.284 µg•mL$^{-1}$ leupeptin, 1.37 µg•mL$^{-1}$ pepstatin A, 0.17 mg•mL$^{-1}$ PMSF, and 0.33 mg•mL$^{-1}$ benzamidine). The amylose column was washed with ten column volumes of lysis buffer, followed by five column volumes of high salt buffer (1000 mM NaCl, 20 mM Na-HEPES pH 7.4, 10% (v/v) glycerol, and 1 mM DTT) and five column volumes lysis buffer. The protein was eluted using amylose elution buffer (300 mM NaCl, 20 mM Na-HEPES pH 7.4, 10% (v/v) glycerol, 116 mM maltose, and 1 mM DTT). Peak fractions were pooled and mixed with 1.5 mg of His$_6$-TEV protease and dialyzed overnight against lysis buffer. The protein was then applied to a HiLoad S200 16/600 pg column equilibrated in SEC buffer (300 mM NaCl, 20 mM Na-HEPES pH 7.4, 10% (v/v) glycerol, and 1 mM DTT). The peak fractions were pooled, concentrated in 10 kDa MWCO Amicon Ultra Centrifugal Filter (Millipore) to 5 mL and then applied to an amylose column equilibrated in SEC buffer to remove the cleaved MBP tag. Protein purity was assessed by SDS-PAGE followed by Coomassie staining. Peak fractions were concentrated in 10 kDa MWCO Amicon Ultra Centrifugal Filter (Millipore). The protein was stored in the SEC buffer. The protein was aliquoted, flash-frozen in liquid nitrogen, and stored at −80°C.

The BRD4-PID constructs were purified as follows: Two liters of *E. coli* Bl21 (DE3) RIL were used for each protein preparation. Cell pellets were thawed, lysed by sonication, and cleared by centrifugation. The clarified lysate was then applied to a HisTrap equilibrated in lysis buffer (300 mM NaCl, 20 mM Na-HEPES pH 7.4, 10% (v/v) glycerol, and 1 mM DTT, 0.284 µg•mL$^{-1}$ leupeptin, 1.37 µg•mL$^{-1}$ pepstatin A, 0.17 mg•mL$^{-1}$ PMSF, and 0.33 mg•mL$^{-1}$ benzamidine). The column was washed with ten column volumes of lysis buffer followed by five column volumes high salt buffer (1000 mM NaCl, 20 mM Na-HEPES pH 7.4, 10% (v/v) glycerol, and 1 mM DTT) and lysis buffer for five column volumes. The column was developed over a 30-minute gradient containing nickel elution buffer (300 mM NaCl, 20 mM Na-HEPES pH 7.4, 10% (v/v) glycerol, 500 mM imidazole pH 8.0, and 1 mM DTT) at a flow rate of 1.5 mL/min. Peak fractions were pooled and mixed with 1.5 mg of His$_6$-TEV protease and dialyzed overnight against lysis buffer. The protein was then applied to a HisTrap equilibrated in lysis buffer to remove the His$_6$-TEV protease, uncleaved protein, and the His$_6$-MBP tag. Protein was concentrated in a 3 kDa MWCO Amicon Ultra Centrifugal Filter (Millipore) and applied to a HiLoad S75 16/600 pg column equilibrated in SEC buffer (300 mM NaCl, 20 mM Na-HEPES pH 7.4, 10% (v/v) glycerol, and 1 mM DTT). Protein purity was assessed by SDS-PAGE and Coomassie staining. Peak fractions were concentrated in a 3 kDa MWCO Amicon Ultra Centrifugal Filter (Millipore). The protein was stored in the SEC buffer. The protein was aliquoted, flash-frozen in liquid nitrogen, and stored at −80°C.

The Avi-BRD4-PID was purified like BRD4-PID except for the following modifications. After eluting from the HisTrap column, proteins were concentrated in 10 kDa MWCO Amicon Ultra Centrifugal Filter (Millipore) to 1 mL and incubated with BirA-$His_6$, 500 µM biotin, 1.0 mg TEV protease, 10 mM ATP pH 7.0, and 10 mM $MgCl_2$ for 2 hours at 30°C. The reaction was incubated at 4°C overnight. The protein was then applied to a HisTrap column in tandem with a HiLoad S75 16/600 pg column equilibrated in SEC buffer.

The AFF4 (1–98) construct was expressed in two liters of *E. coli* Rosetta 2 (DE3) pLysS. Cell pellets were thawed, lysed by sonication, and cleared by centrifugation. The clarified lysate was then applied to a 5 mL HisTrap column equilibrated in lysis buffer (500 mM NaCl, 20 mM Na-HEPES pH 7.4, 10% (v/v) glycerol, and 1 mM DTT, 0.284 µg•$mL^{-1}$ leupeptin, 1.37 µg•$mL^{-1}$ pepstatin A, 0.17 mg•$mL^{-1}$ PMSF, and 0.33 mg•$mL^{-1}$ benzamidine). The column was washed with ten column volumes of lysis buffer followed by five column volumes high salt buffer (1000 mM NaCl, 20 mM Na-HEPES pH 7.4, 10% (v/v) glycerol, and 1 mM DTT) and five column volumes lysis buffer. The column was developed over a 30-minute gradient using nickel elution buffer (500 mM NaCl, 20 mM Na-HEPES pH 7.4, 10% (v/v) glycerol, 500 mM imidazole pH 8.0, and 1 mM DTT). Peak fractions were pooled and mixed with 1.5 mg of $His_6$-TEV protease and dialyzed overnight against lysis buffer. The protein was then applied to a HisTrap column equilibrated in lysis buffer to remove the $His_6$-TEV protease, uncleaved protein, and the $His_6$-tag. Protein was concentrated in a 3 kDa MWCO Amicon Ultra Centrifugal Filter (Millipore) and applied to a HiLoad S75 16/600 pg column equilibrated in SEC buffer (300 mM NaCl, 20 mM Na-HEPES pH 7.4, 10% (v/v) glycerol, and 1 mM DTT). Protein purity was assessed by SDS-PAGE followed by Coomassie staining. Peak fractions were concentrated in 3 kDa MWCO Amicon Ultra Centrifugal Filters (Millipore). The protein was aliquoted, flash-frozen in liquid nitrogen, and stored at −80°C.

HEXIM1 was purified from four liters of *E. coli* Rosetta 2 (DE3) pLysS. Cell pellets were thawed and lysed by sonication. After sonication, polyethylenimine was added to the lysate to a final concentration of 0.2% (w/v) while the lysate was stirring. The lysate was left to incubate for five minutes while stirring before it was cleared by centrifugation. The clarified lysate was then applied to a 5 mL HisTrap column equilibrated in lysis buffer (300 mM NaCl, 20 mM Na-HEPES pH 7.4, 10% (v/v) glycerol, and 1 mM DTT, 0.284 µg•$mL^{-1}$ leupeptin, 1.37 µg•$mL^{-1}$ pepstatin A, 0.17 mg•$mL^{-1}$ PMSF, and 0.33 mg•$mL^{-1}$ benzamidine). The column was washed with ten column volumes of lysis buffer followed by five column volumes high salt buffer (1000 mM NaCl, 20 mM Na-HEPES pH 7.4, 10% (v/v) glycerol, and 1 mM DTT) and lysis buffer for five column volumes. The column was developed over a 30-minute gradient against nickel elution buffer (300 mM NaCl, 20 mM Na-HEPES pH 7.4, 10% (v/v) glycerol, 500 mM imidazole pH 8.0, and 1 mM DTT). Peak fractions were pooled and mixed with 1.5 mg of $His_6$-TEV protease and dialyzed overnight against lysis buffer. The protein was then applied to a HisTrap equilibrated in lysis buffer to remove the $His_6$-TEV protease, uncleaved protein, and the $His_6$ tag. Protein was concentrated in a 30 kDa MWCO Amicon Ultra Centrifugal Filter (Millipore) and applied to a HiLoad S200 16/600 pg column equilibrated in SEC buffer (300 mM NaCl, 20 mM Na-HEPES pH 7.4, 10% (v/v) glycerol, and 1 mM DTT). Protein purity was assessed by SDS-PAGE followed by Coomassie staining. Peak fractions were concentrated in 30 kDa MWCO Amicon Ultra Centrifugal Filter (Millipore). The protein was stored in the SEC buffer. The protein was aliquoted, flash-frozen in liquid nitrogen, and stored at −80°C.

The RPB1-CTD construct was purified from six liters of *E. coli* Bl21 (DE3) RIL. Cell pellets were thawed, lysed by sonication, and cleared by centrifugation. The clarified lysate was applied to a 5 mL HisTrap column equilibrated in lysis buffer (300 mM NaCl, 20 mM Na-HEPES pH 7.4, 10% (v/v) glycerol, and 1 mM DTT, 0.284 µg•$mL^{-1}$ leupeptin, 1.37 µg•$mL^{-1}$ pepstatin A, 0.17 mg•$mL^{-1}$ PMSF, and 0.33 mg•$mL^{-1}$ benzamidine). The column was washed with ten column volumes of lysis buffer followed by five column volumes high salt buffer (1000 mM NaCl, 20 mM Na-HEPES pH 7.4, 10% (v/v) glycerol, and 1 mM DTT) and lysis buffer for five column volumes. A column packed with 10 mL of amylose resin equilibrated in lysis buffer was attached to the base of the HisTrap column. The HisTrap column was developed over a 30-minute gradient against nickel elution buffer (300 mM NaCl, 20 mM Na-HEPES pH 7.4, 10% (v/v) glycerol, 500 mM imidazole pH 8.0, and 1 mM DTT). The HisTrap column was removed, and the amylose column was washed with five column volumes of lysis buffer. The protein was eluted using amylose elution buffer (300 mM NaCl, 20 mM Na-HEPES pH 7.4, 10% (v/v) glycerol, 116 mM maltose, and 1 mM DTT). Peak fractions were concentrated in a 30 kDa MWCO Amicon Ultra Centrifugal Filter (Millipore). The protein was then applied to a HiLoad S200 16/600 pg column equilibrated in SEC buffer (300 mM NaCl, 20 mM Na-HEPES pH 7.4, 10% (v/v) glycerol, and 1 mM DTT). Protein purity was assessed by SDS-PAGE followed by Coomassie staining. Peak fractions were concentrated in a 30 kDa MWCO Amicon Ultra Centrifugal Filter (Millipore). The protein was stored in the SEC buffer. The protein was aliquoted, flash-frozen in liquid nitrogen, and stored at −80°C.

#### Preparation of Alexa Fluor 555 labeled WT BRD4-PID

Purified WT BRD4-PID containing an N-terminal cysteine residue was dialyzed overnight at 4°C against dialysis buffer containing 300 mM NaCl, 20 mM Na-HEPES pH 7.4, 10% (v/v) glycerol, and 1 mM TCEP pH 7.0 in a 3.5 kDa Slide-A-Lyzer mini dialysis unit to remove DTT. Alexa Fluor 555 C2-maleimide (ThermoFisher) was resuspended in dialysis buffer to a concentration of 10 mM. Cysteine bearing WT BRD4-PID was mixed with a 10-fold molar excess of Alexa Fluor 555 C2-maleimide and incubated overnight at room temperature on a rotating wheel. The labeling reaction was quenched by adding DTT to a final concentration of 10 mM. Excess dye was removed by applying the protein to a G-10 desalting spin column equilibrated in buffer (Cytiva) followed by multiple rounds of diafiltration in a 3 kDa MWCO Amicon Ultra 0.5 mL filter. Labelling efficiency was determined by measuring the signal at $A_{555}$ nm and dividing it by the concentration of the protein and multiplying by 150,000. The labeling efficiency was determined to be 17.3%.

#### Analytical Size-Exclusion Chromatography

P-TEFb (5 µM) and BRD4-PID (10 µM) were incubated in a final buffer containing 100 mM NaCl, 20 mM Na-HEPES pH 7.4, 10% (v/v) glycerol, and 1 mM DTT and incubated for 30 minutes at 30°C. The complex was applied to a Superdex 200 3.2/300 column equilibrated in complex buffer on an Äkta Micro purification system (Cytiva) at 4 °C (100 mM NaCl, 20 mM Na-HEPES pH 7.4, 4% (v/v) glycerol, 3 mM $MgCl_2$, and 1 mM DTT). Additionally, P-TEFb and BRD4-PID were applied to a Superdex 200 3.2/300 column equilibrated in complex buffer on an Äkta Micro purification system (Cytiva) at 4 °C on their own. Peak fractions were analyzed by SDS-PAGE using a 4–12% gradient gel followed by Coomassie staining.

#### Biolayer interferometry

All protein used for biolayer interferometry (BLI) were dialyzed against 500 mL of BLI assay buffer (20 mM Na-HEPES pH 7.4, 100 mM NaCl, 0.02% Tween-20, 1 mM DTT) for at least four hours at 4°C. Octet SA biosensors (Sartorius) coated with streptavidin were incubated in BLI assay buffer for 10 minutes at 30 °C. The SA biosensors were incubated with BLI buffer for 60 seconds, then moved to wells of 200 µL of 10 nM of Avi-BRD4-PID for 300 seconds. The SA biosensors were then incubated with serially diluted P-TEFb (WT) or P-TEFb (C-lobe mutant) (Dilution series 0-1000 nM) for 300 seconds to measure association. The SA biosensors were then incubated with BLI buffer for 600 seconds to measure dissociation for the Avi-BRD4-PID construct. Throughout the experiment the plate was shaking at 1000 rpm and kept at 30 °C. A set of SA biosensors that were not incubated with Avi-tagged protein were also applied to the P-TEFb titration series to account for nonspecific binding. The signal was subtracted from the no Avi-tagged protein and the 0 nM P-TEFb control for the association and dissociation curve. The experiments were performed in triplicate. Binding curves were fit for each replicate using the 333 nM, 111 nM, and 37 nM concentrations because these had $r^2$ fits of ≥0.9, in GraphPad Prism Version 10 and the average and standard deviation of the $K_d$ from each binding replicate were determined and reported.

#### NADH-coupled ATPase Assay

A modified NADH coupled ATPase assay was used to measure relative rates of ATP hydrolysis of P-TEFb in the presence of BRD4-PID as previously described.[1,3] P-TEFb was kept at 0.3 µM in a final buffer containing 3 mM $MgCl_2$, 0.1 mM NADH, 0.4% (w/v) pyruvate kinase/lactate dehydrogenase, 1 mM phosphoenolpyruvate, 100 mM NaCl, 20 mM Na-HEPES pH 7.4, 10% (v/v) glycerol and 1 mM DTT. BRD4-PID was titrated (0–10 µM). 5 µM of MBP-RPB1-CTD was used. Samples were incubated for 5 min at 30 °C before adding 1 mM ATP pH 7.0. The absorption at 340 nm was measured in 384-well, clear flat bottom, square well plates (Greiner: 781101) over 60 min at 30 °C in a Tecan Spark plate reader. The rate of change in absorbance at 340 nm over time was determined from the linear region of the resulting absorbance curves. All experiments were performed three times.

For the 7SK release assays, experiments were performed as above except that complexes containing P-TEFb, P-TEFb and the 7SK RNA, P-TEFb and HEXIM1, and P-TEFb, the 7SK RNA, and HEXIM1, were first incubated in a buffer containing 100 mM NaCl, 20 mM Na-HEPES pH 7.4, 10% (v/v) glycerol, 3 mM $MgCl_2$, and 1 mM DTT for 30 minutes at 30 °C, before being used in the assays at a final concentration of 0.3 µM. BRD4-PID and AFF4 (1–98) were titrated (0–10 µM). The 7SK RNA was synthesized by IDT and contained a FAM label on the 5' end of the RNA and encompassed residues 26–85 of the 7SK RNA: RNA (5'-6-FAM-CAU CUG UCA CCC CAU UGA UCG CCA GGG UUG AUU CGG CUG AUC UGG CUG GCU AGG CGG UG-3').

#### Gel-based P-TEFb Activity Assay

P-TEFb (0.1 µM) was incubated with 10 µM BRD4-PID and 5 µM of MBP-RPB1-CTD in a final buffer containing 3 mM $MgCl_2$, 100 mM NaCl, 20 mM Na-HEPES pH 7.4, 10% (v/v) glycerol and 1 mM DTT for 5 minutes at 30 °C before adding 1 mM ATP pH 7.0 in a final volume of 50 µL. 5 µL of the reaction was removed after 0, 1, 5, 15, 30, 60, and 120 minutes after ATP addition and mixed with 5 µL of 2x SDS loading dye buffer (100 mM Tris-HCl pH 6.8, 4% (w/v) SDS, 20% (v/v) glycerol, 0.01% (w/v) bromophenol blue, 10 mM 2-mercaptoethanol). This experiment was also done with 0.1 µM P-TEFb incubated with 5 µM of MBP-RPB1-CTD and with 10 µM BRD4-PID incubated with 5 µM of MBP-RPB1-CTD as controls. Time points were analyzed by SDS-PAGE using 4–12% gradient SDS-PAGE gels followed by Coomassie staining. Experiments were performed three times.

#### AlphaFold Modeling

AlphaFold 3 modeling was performed on the online AlphaFold server.[4] For the P-TEFb–BRD4-PID AlphaFold 3 model CDK9 residues 1–330 were co-folded with CYCT1 residues 1–259 and BRD4 residues 1300–1362. For the P-TEFb–HEXIM1 AlphaFold 3 model, two molecules of CDK9 residues 1–372 were co-folded with two molecules of CYCT-1 residues 1–272 and two molecules of HEXIM1 residues 1–359. Models were visualized in ChimeraX v1.10.1.[5]

#### Cryo-EM Sample Preparation

P-TEFb (4 µM) was incubated with 20 µM BRD4-PID and 2 mM ATPγS in a final buffer of 100 mM NaCl, 20 mM Na-HEPES pH 7.4, 3.5% (v/v) glycerol, 6 mM $MgCl_2$, and 1 mM DTT. The sample was incubated for 30 minutes at 30 °C shaking at 300 rpm. The sample was applied to R2/2 UltrAuFoil grids (Quantifoil) that were glow discharged for 2 minutes using a K100X glow discharger before 2 µL of sample was applied to each side of the grid. The sample was incubated for 4 seconds and blotted for 1 second. Vitrification was done by plunging the grids into liquid ethane using a Vitrobot Mark IV (FEI Company) maintained at 5 °C and 100% humidity.

#### Cryo-EM Data Collection and Processing

Cryo-EM data were collected on a FEI Titan Krios operated at 300 kV. A K3 summit direct detector with a GIF quantum energy filter (Gatan) was operated with a slit width of 20 eV. Automated data acquisition was performed with FEI EPU v2.12.1 software at ominal magnification of 130,000x, corresponding to a pixel size of 0.654 Å/px. Data acquisition was done with a defocus range of 0.2–1.4 µm. Image stacks of 70 frames were collected over 1 s in counting mode. The total dose for the sample was 70.0 $e^-/Å^2$. A total of 11,484 image stacks were collected. A second data collection from the same grid was done at a 15º tilt to help combat orientation bias. Data acquisition was done with a defocus range of 0.2–1.4 µm. Image stacks of 71 frames were collected over 1 s in counting mode. The total dose for the sample was 70.65 $e^-/Å^2$. A total of 14,087 image stacks were collected.

Each data set was initially separately processed in cryoSPARC v4.7, with later processing in cryoSPARC v5.0.2.[6] The image stacks were motion corrected and the contrast transfer function was estimated. Particles were picked using two blob pickers: an elliptical blob picker from 50–150 Å and a circular blob picker from 80–110 Å. Particles were extracted at a box size of 352px and Fourier cropped to 176px. Each particle set was individually subjected to 2D classification with the following parameters (all other parameters were left at their default values): batch size per class = 5000, Number of online EM iterations = 50, Force max over poses/shifts = off, Initial uncertainty factor = 1, number of 2D classes = 100, circular mask diameter = 100 Å. Each particle set underwent 2 rounds of 2D classification, removing 2D classes that represented junk classes after every round of classification. Particles were then re-extracted to re-center them and duplicate particles within each data set were removed. Another round of 2D classification was done, and the junk classes were removed. The particles from both data sets were then combined and subjected to two final rounds of 2D classification.

After this final round of 2D classification, the particle stack contained 5,847,708 particles. All particles were subjected to an ab-initio reconstruction with the following parameters (all other parameters were left at their default values): number of ab-initio classes = 10, maximum resolution = 9, initial resolution = 12, initial minibatch size = 300, final minibatch size = 1000, class similarity = 0. on-uniform refinement job was performed with the particles and volume of the ab-initio reconstruction class that best resembled P-TEFb and that had extra density for the BRD4-PID, which contained 887,702 particles, with the volume as an input low-pass filtered to 12 Å. The refined volume was then subjected to a local refinement with the following parameters (all other parameters were left at their default values): re-center rotations each iteration = true, re-center shifts each iteration = true, force re-do GS split = true, minimize over per-particle scale = true, rotation search ext = 2 deg, shift search ext = 1 Å). The local refinement was done with the output mask from the non-uniform refinement, which encompassed the whole complex. The locally refined volume was then subjected to 3D classification with the following parameters (all other parameters were left at their default values): number of classes = 2, filter resolution = 3 Å, batch size per class = 5000, initial structure low pass filter = 12 Å, force hard classification = true, initialization mode = PCA, number of O-EM epochs = 5. The most occupied class contained 475,526 particles and was subjected to local refinement using the parameters above and the particles were re-extracted at a box size of 352 px. The particles were then used in a local refinement with the following parameters (all other parameters were left at their default values): re-center rotations each iteration = true, re-center shifts each iteration = true, force re-do GS split = true, minimize over per-particle scale = true, rotation search ext = 2 deg, shift search ext = 1 Å. The particle stack was then subjected to another round of 3D classification using the following parameters (all other parameters were left at their default values): number of classes = 2, filter resolution = 3 Å, batch size per class = 5000, initial structure low pass filter = 12 Å, initialization mode = PCA, number of O-EM epochs = 5, Use latent mixing coefficients = true. The most occupied class contained 352,345 particles and was locally refined using the parameters above and then used in 3D classification using the following parameters (all other parameters were left at their default values): number of classes = 2, filter resolution = 3 Å, batch size per class = 5000, initial structure low pass filter = 20 Å, initialization mode = PCA, number of O-EM epochs = 5, Use latent mixing coefficients = true. The most occupied class contained 238,413 particles and was subjected to another local refinement followed by reference-based motion correction and another round of local refinement. The particle stack was then imported into Relion-5.[7] 3D auto-refine was performed on the particle stack using the following parameters (all other parameters were left at their default values): Initial low pass filter = 15 Å, Mask diameter = 200 Å, Blush regularization = ON, Initial angular sampling = 3.7 degrees, Local searches from auto-sampling = 1.8 degrees, resulting in Map 1. Map 1 was then sharpened using a b-factor of –245 and used for model building, resulting in Map 2.

#### Model Building

Each chain of an AlphaFold 3 model of the P-TEFb–BRD4-PID complex was rigid body fit into Map 2 using ChimeraX v1.10.1.[5] To improve the model fit and to remove low confidence parts of the model, the model was subjected to Phenix v1.20.1-4487 dock_and_rebuild.[8] The model was then flexibly fit into map 2 using the online Namdinator server using default parameters at an expected resolution of 5 Å.[9] ATPγS was modelled by aligning a previously determined structure of human CDK7 bound to ATPγS (PDB 8P6Y) onto our model, extracting the coordinates for ATPγS from this model, and integrating the ATPγS coordinates into our model.[10] The model was manually adjusted in Coot v1.3.1 and was real-space refined in Phenix v1.20.1-4487.[8,11] ReadySet was used to prepare input PDB models for refinement. Real space refinement was performed with global minimization, occupancy, nqh flips, local grid search, and adp on, with max iterations of 1000 and 10 macro cycles, and unbonded weight of 1000. Data collection and modeling statistics are in Table S1.

#### Native PAGE with Alexa-Fluor 555 labeled BRD4-PID

P-TEFb (1.0 μM) was incubated with 0.05 μM Alexa Fluor 555 labeled BRD4-PID. Either unlabeled BRD4-PID, AFF4 (1–98), BRD4-PID (LLAA), or BRD4-PID (Helix-1 mutant) was titrated (0–10 μM) into the samples containing P-TEFb and labeled BRD4-PID and were then incubated on ice for 10 minutes. Samples were then incubated in a final buffer containing 100 mM NaCl, 20 mM Na-HEPES pH 7.4, 10% (v/v) glycerol and 1 mM DTT for 10 minutes at room temperature. Samples were loaded on a 4–16% Native PAGE gel

(Invitrogen) that was pre-run for 45 minutes in 1x Native PAGE buffer (Invitrogen) at 150 V at 4°C. Samples were run for 120 minutes at the same voltage and imaged using a Typhoon 9500 FLA Imager (GE Healthcare Life Sciences) at a PMT of 1000 to detect the Alexa Fluor 555 label. All experiments were done in triplicate. Gels were imported into Fiji (v2.1.0). The band intensity signal for the free labeled BRD4-PID and the bound labeled BRD4-PID for each lane were measured. A box for the free labeled BRD4-PID with an area of 7650 pixels$^2$ and a box for the bound labeled BRD4-PID with an area of 16443 pixels$^2$ were used for the quantification. The bands were background subtracted and the ratio of the signal at the free band was taken to the additive signal of the free and bound bands. This was done for all gels, and the data was plotted using GraphPad Prism.

#### Multiple Sequence Alignment

Sequences for CDK9, BRD4, and AFF4 were obtained from UniProt. Sequences were aligned using MAFFT v7 online server and visualized in JalView v2.11.5.1.[12,13]

#### Fluorescence Anisotropy

The SL1 7SK RNA was synthesized by IDT and contained a FAM label on the 5' end of the RNA and encompassed residues 26–85 of the 7SK RNA: RNA (5'-6-FAM-CAU CUG UCA CCC CAU UGA UCG CCA GGG UUG AUU CGG CUG AUC UGG CUG GCU AGG CGG UG-3'). This RNA was incubated at 0.005 µM with either HEXIM1, P-TEFb, BRD4-PID, or AFF4 (1-98). The maximum concentration of the protein used in the assay was 10 µM and each protein was serially diluted. The protein and RNA were incubated together for 10 minutes at room temperature before being supplemented with buffer to bring the final buffer conditions of the assay to 100 mM NaCl, 20 mM Na-HEPES pH 7.4, 10% (v/v) glycerol and 1 mM DTT. The samples were then incubated at room temperature for 10 minutes before being transferred to a Greiner 384 Flat Black Bottom plate. The samples were measured using a Tecan Spark Plate reader using an excitation wavelength of 470 nm and an excitation bandwidth of 20 nm, an emission wavelength of 518 nm and an emission bandwidth of 20 nm, and a gain of 130. The RNA only condition was substracted from the data and plotted using GraphPad Prism. All experiments were done in triplicate.

For the competition binding assays, 0.3 µM of P-TEFb and 0.3 µM of HEXIM1 were incubated with 0.005 µM of the labeled 7SK RNA. Either BRD4-PID or AFF4 (1–98) was serially diluted and added into the sample at final concentrations of 0-10 µM in a final buffer of 100 mM NaCl, 20 mM Na-HEPES pH 7.4, 10% (v/v) glycerol and 1 mM DTT. The samples were incubated at room temperature for 10 minutes before being transferred to a Greiner 384 Flat Black Bottom plate. The samples were measured using a Tecan Spark Plate reader. All experiments were done in triplicate. The data was fitted in GraphPad Prism using the log inhibitor vs response function and the calculated $IC_{50}$ values are reported.

### SUPPLEMENTARY FIGURES

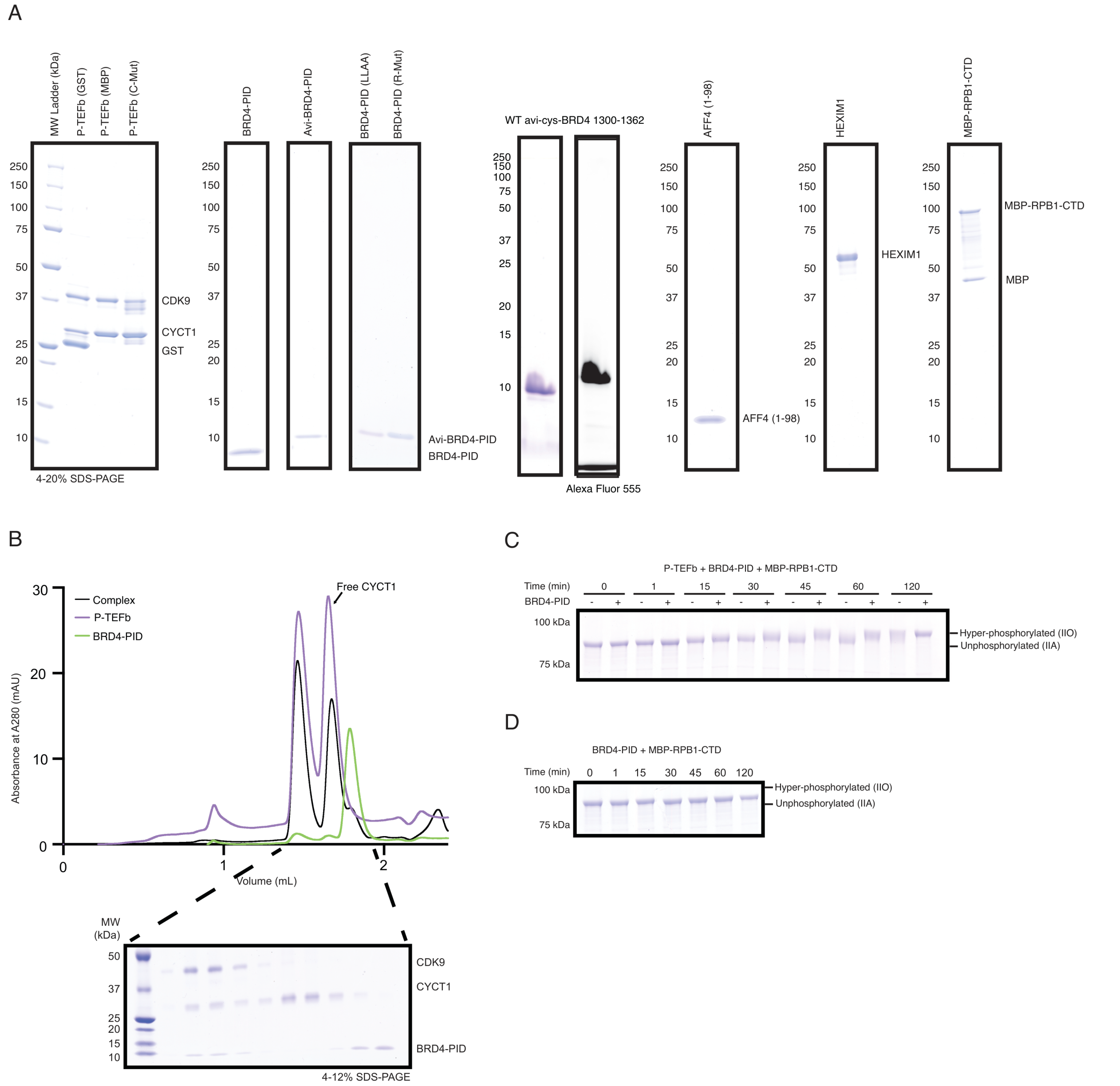

**Figure S1.** Proteins used in this study and biochemical assays characterizing P-TEFb–BRD4-PID interaction. (A) Coomassie stained gels of purified proteins used in this study. 5 µM of protein loaded per well. Molecular weight markers are shown. (B) Analytical size exclusion chromatography traces of the P-TEFb–BRD4-PID complex (black), P-TEFb alone (purple), and BRD4-PID alone (green). Coomassie stained gel of the peak fractions of the complex shown. Molecular weight markers are shown. (C) Full-time course titration of P-TEFb stimulation of RPB1-CTD phosphorylation by BRD4-PID. Coomassie stained gel showing the differential migration of MBP-RPB1-CTD as P-TEFb phosphorylates the protein. Molecular weight markers are shown. The experiment was performed three times. (D) Full-time course titration of BRD4-PID incubated with RPB1-CTD. Coomassie stained gel showing that the migration of MBP-RPB1-CTD does not change with the incubation of only WT BRD4-PID. Molecular weight markers are shown. The experiment was performed 3 times.

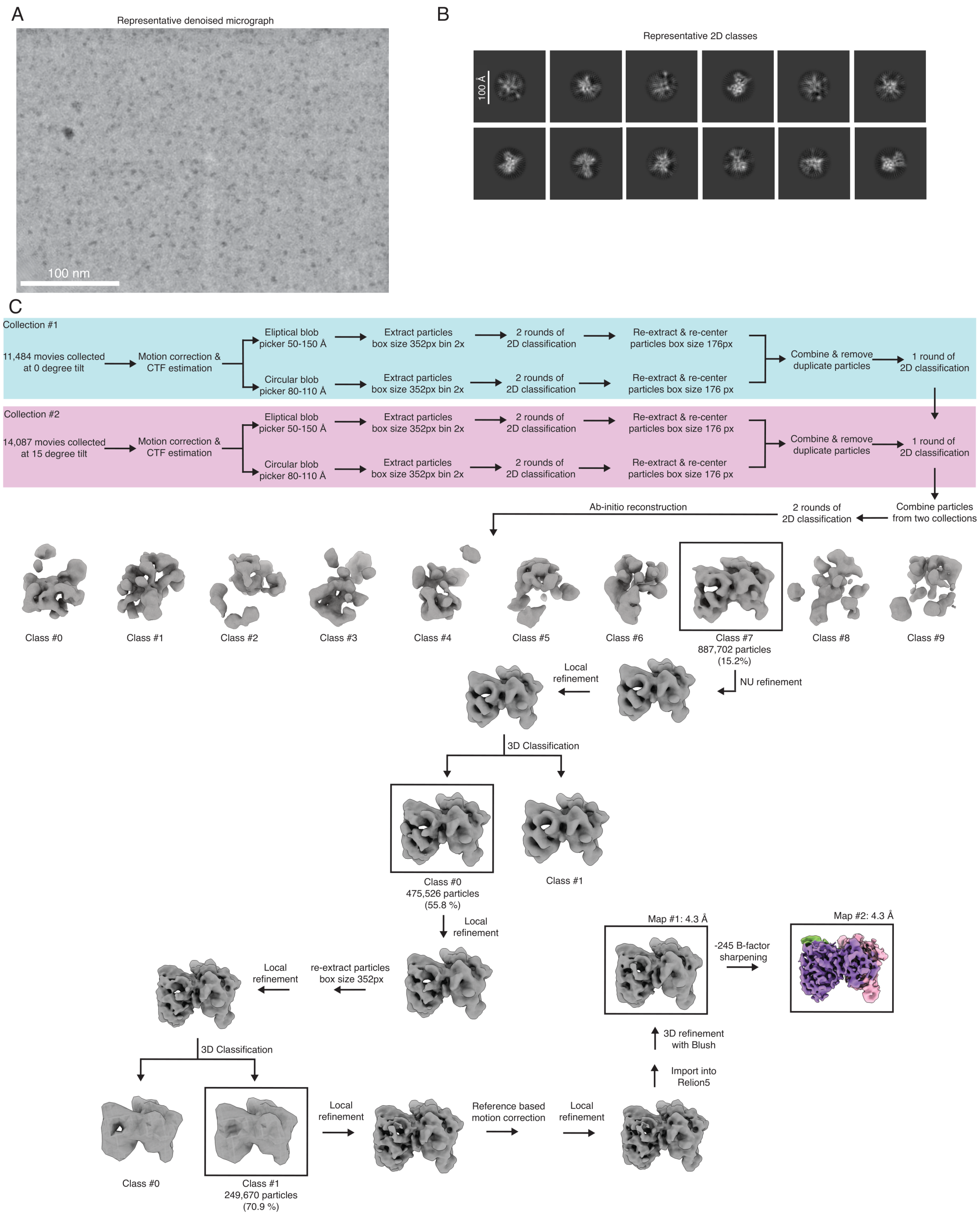

Figure S2. Cryo-EM data processing. (A) Representative denoised micrograph. (B) Representative 2D classes. (C) Cryo-EM data processing tree for the P-TEFb–BRD4-PID complex.

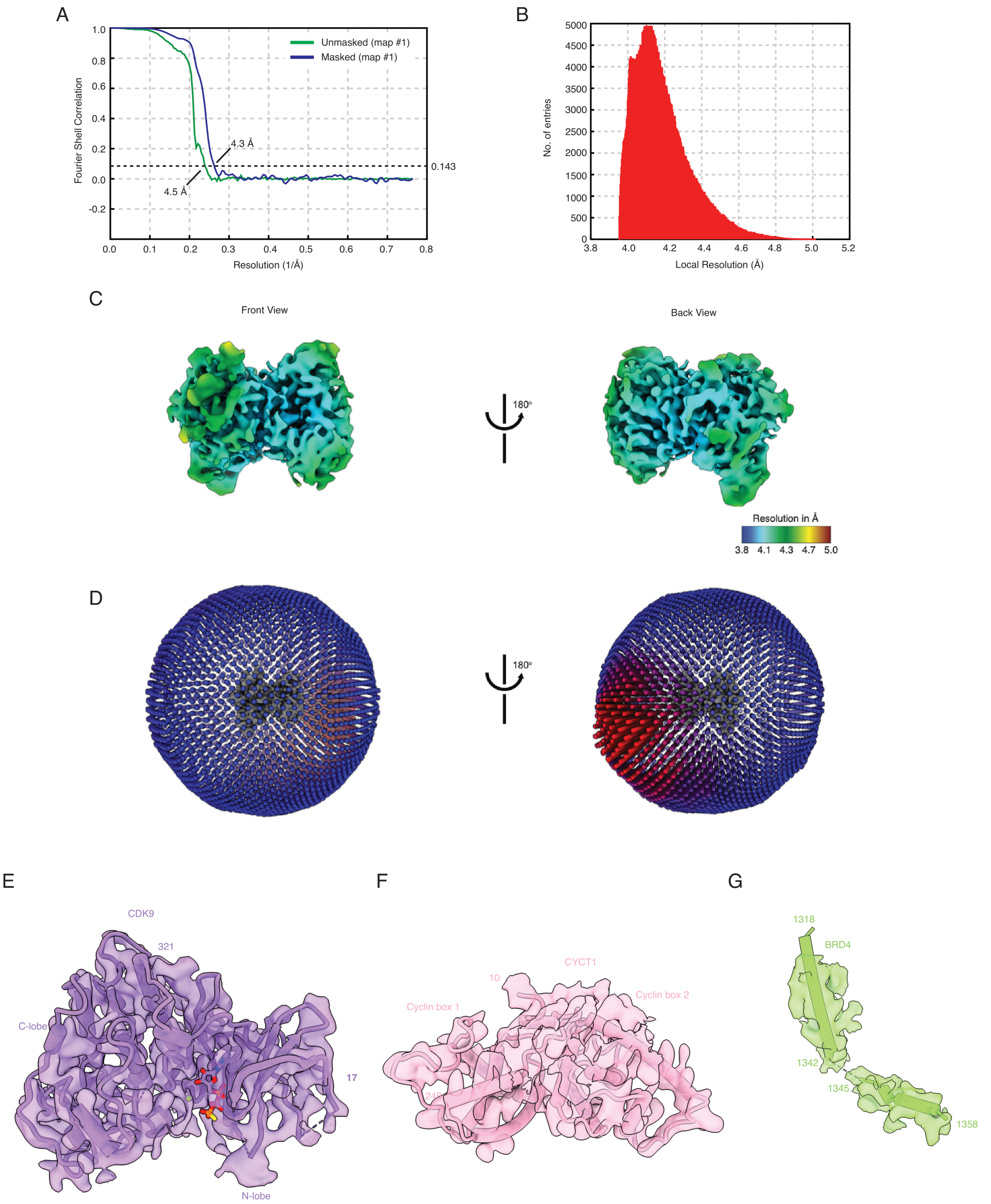

Figure S3. Cryo-EM data statistics. (A) Fourier Shell Correlation (FSC) plots for masked and unmasked maps. (B) Histogram of local resolution. (C) Front and back views of map 2 colored by local resolution. (D) Front and back views of map 2 with orientation distributions. (E) Representative density and model for CDK9 (map 2). (F) Representative density and model for CYCT1 (map 2). (G) Representative density and model for BRD4-PID (map 2).

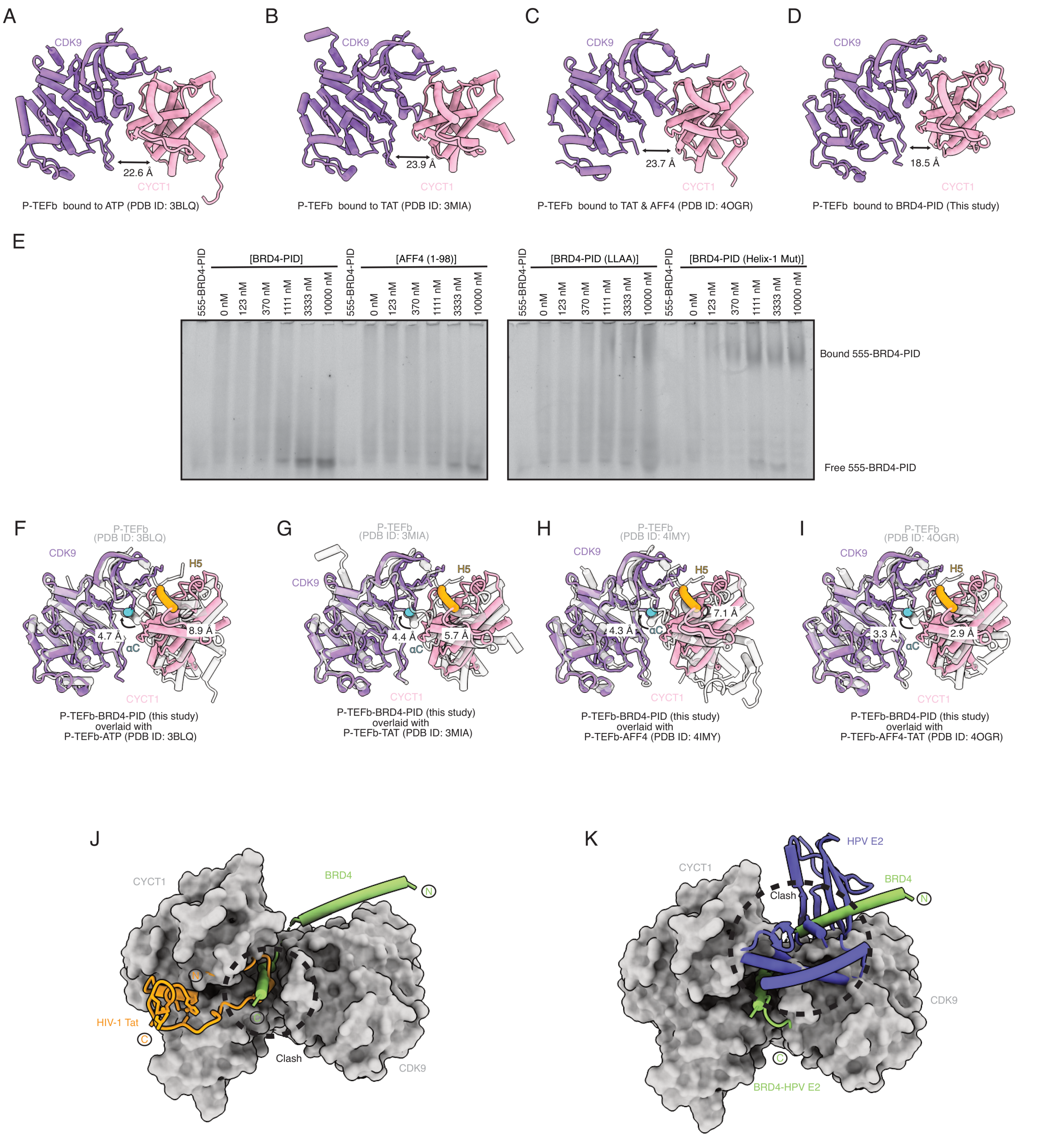

**Figure S4.** Comparison of P-TEFb structures and mutant binding data. (A) Model of P-TEFb bound to ATP. CDK9 is shown in purple and CYCT1 is shown in pink. Measurement between CDK9 (residue 175) and CYCT1 (residue 163) shown. (B) Model of P-TEFb bound to Tat. CDK9 is shown in purple and CYCT1 is shown in pink. Measurement between CDK9 (residue 175) and CYCT1 (residue 163) shown. (C) Model of P-TEFb bound to AFF4 and Tat. CDK9 is shown in purple and CYCT1 is shown in pink. Measurement between CDK9 (residue 175) and CYCT1 (residue 163) shown. (D) Model of P-TEFb bound to BRD4-PID. CDK9 is shown in purple and CYCT1 is shown in pink. Measurement between CDK9 (residue 175) and CYCT1 (residue 163) shown. (E) Representative native gel assays used for quantifications in Figure 3D and Figure 4C. (F–I) Complete structural alignment of P-TEFb–BRD4 to (F) P-TEFb bound to ATP (PDB ID: 3BLQ), (G) P-TEFb bound to Tat (PDB ID: 3MIA), (H) P-TEFb bound to AFF4 (PDB ID: 4IMY), and (I) P-TEFb bound to AFF4–HIV-1 Tat (PDB ID: 4OGR). Only CDK9 and CYCT1 are shown for clarity with CDK9 (purple) and CYCT1 (pink) from the P-TEFb–BRD4 model. All other P-TEFb models shown in gray. CDK9 αC shown in blue and CYCT1 H5 shown in yellow. Measurements as in Figure 3. (J) HIV-1 Tat (orange) (PDB ID: 3MIA)

modeled onto P-TEFb (gray in surface) bound to BRD4-PID (green) model. Clash between Tat and BRD4-PID indicated. (K) HPV E2 (blue) (PDB ID: 2NNU) modeled onto the P-TEFb (gray in surface) bound to BRD4-PID (green) model. Clash between E2 and BRD4-PID is indicated.

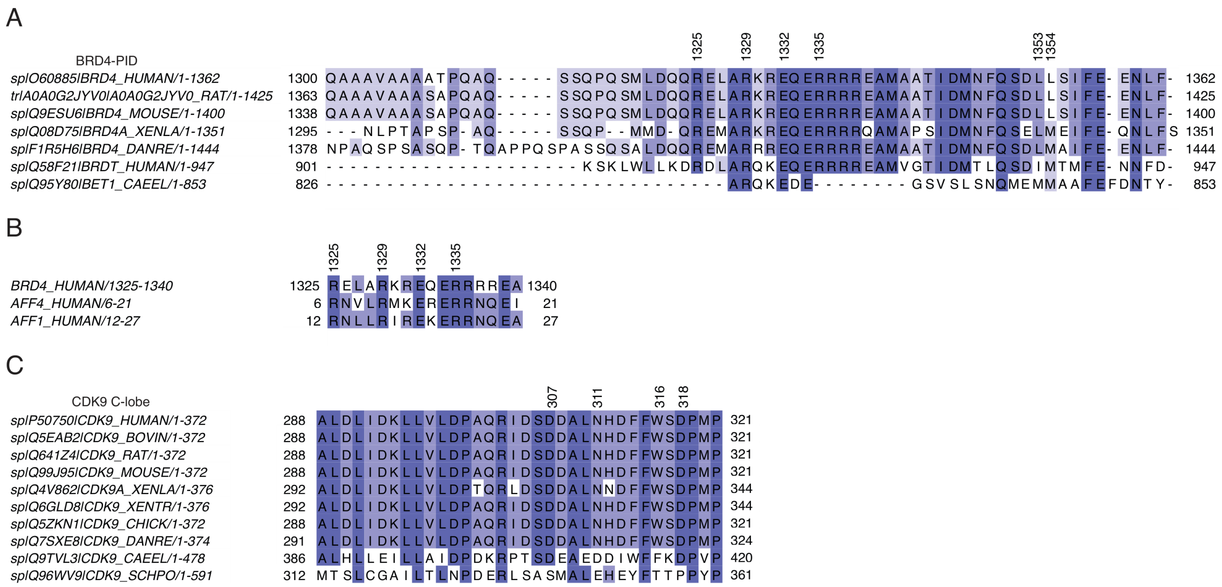

Figure S5. Sequence Alignments for CDK9, AFF4, AFF1, and BRD4. (A) Sequence alignment for BRD4 residues 1300–1362. (B) Sequence alignment for BRD4 residues 1325–1340, AFF4 residues 6–21, and AFF1 residues 12–27. (C) Sequence alignment for CDK9 residues 288–321.

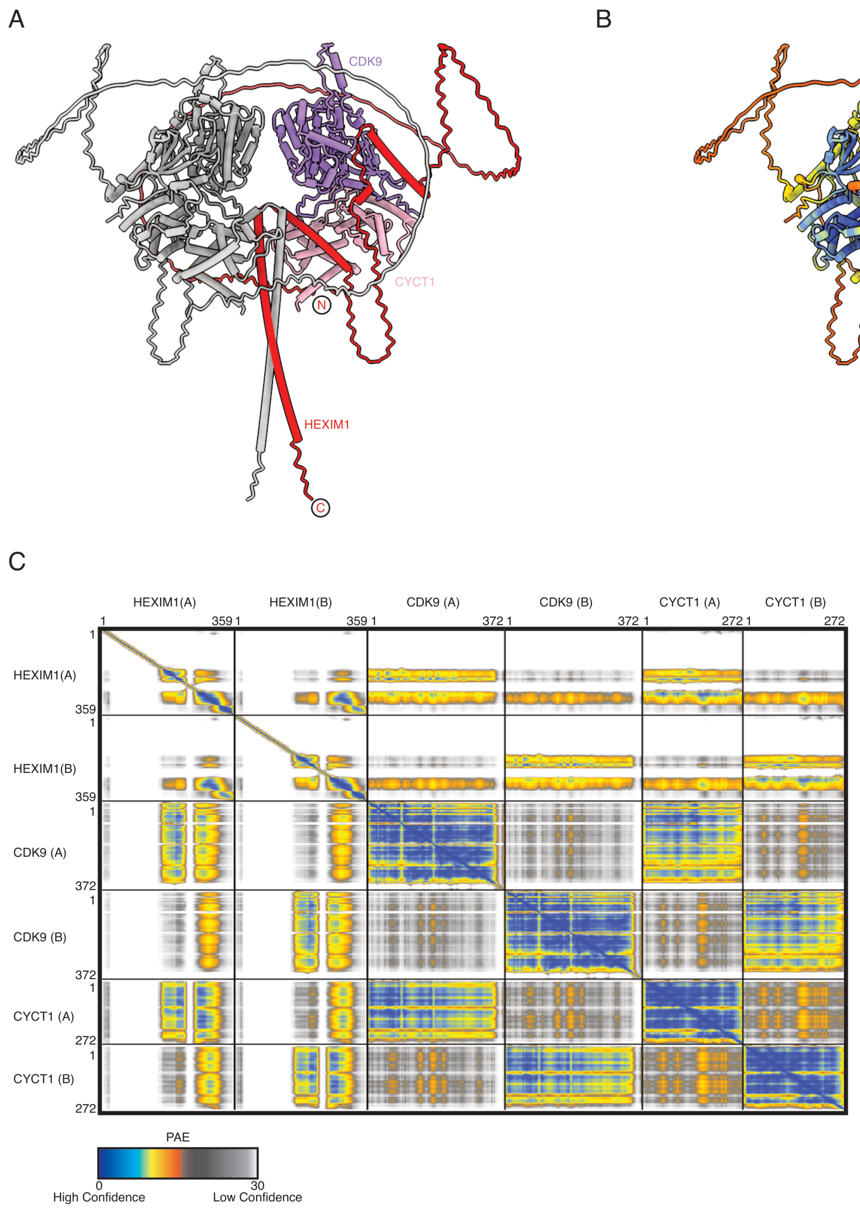

Figure S6. AlphaFold 3 modeling for HEXIM1–P-TEFb complex. (A) P-TEFb–HEXIM1 dimer shown with one half of the dimer colored in gray and the other half colored with CDK9 in purple, CYCT1 in pink, and HEXIM1 in red. (B) P-TEFb–HEXIM1 dimer colored with the pLDDT scores, scale bar shown. (C) P-TEFb–HEXIM1 prediction PAE plot, scale bar shown.

#### SUPPLEMENTARY TABLE

**Table S1. Cryo-EM data collection, refinement, and validation statistics.**

| | **P-TEFb–BRD4-PID Complex (EMD-XXXX) (PDB: XXXX)** |
|---|---|
| Data collection and processing | |
| Magnification | 130,000 |
| Voltage (kV) | 300 |
| Electron exposure (e–/Å$^2$) | 70.0, 70.65 |
| Defocus range (μm) | 0.2−1.4 |
| Pixel size (Å) | 0.654 |
| Symmetry imposed | C1 |
| Initial particle images (no.) | 5,847,708 |
| Final particle images (no.) | 238,413 |
| Map resolution (Å) | 4.3 |
| FSC threshold | 0.143 |
| Map resolution range (Å) | 4.0−5.0 |
| Refinement | |
| Initial models used (PDB code) | AlphaFold 3 Model<br>8P6Y |
| Map sharpening B factor (Å$^2$) | −245 |
| Model composition | |
| Non-hydrogen atoms | 4616 |
| Protein residues | 562 |
| Nucleotides | 0 |
| Ligands | AGS: 1 |
| B factors (Å2) | |
| Protein | 73.15 |
| Nucleotide | 0 |
| Ligand | 75.37 |
| R.m.s. deviations | |
| Bond lengths (Å) | 0.004 |
| Bond angles (°) | 0.888 |
| Validation | |
| MolProbity score | 1.71 |
| Clashscore | 7.02 |
| Poor rotamers (%) | 0.00 |
| Ramachandran plot | |
| Favored (%) | 95.42 |
| Allowed (%) | 4.58 |
| Disallowed (%) | 0.00 |
